# Peripheral Coupling Sites Formed by STIM1 Govern the Contractility of Vascular Smooth Muscle Cells

**DOI:** 10.1101/2021.05.25.445565

**Authors:** Vivek Krishnan, Sher Ali, Albert L. Gonzales, Pratish Thakore, Caoimhin S. Griffin, Evan Yamasaki, Michael G. Alvarado, Martin T. Johnson, Mohamed Trebak, Scott Earley

## Abstract

Peripheral coupling between the sarcoplasmic reticulum (SR) and plasma membrane (PM) forms signaling complexes that regulate the membrane potential and contractility of vascular smooth muscle cells (VSMCs), although the mechanisms responsible for these membrane interactions are poorly understood. In many cells, STIM1 (stromal-interaction molecule 1), a single transmembrane-domain protein that resides in the endoplasmic reticulum (ER), transiently moves to ER-PM junctions in response to depletion of ER Ca^2+^ stores and initiates store-operated Ca^2+^ entry (SOCE). Fully differentiated VSMCs express STIM1 but exhibit only marginal SOCE activity. We hypothesized that STIM1 is constitutively active in contractile VSMCs and maintains peripheral coupling. In support of this concept, we found that the number and size of SR-PM interacting sites were decreased and SR-dependent Ca^2+^ signaling processes were disrupted in freshly isolated cerebral artery SMCs from tamoxifen-inducible, SMC-specific STIM1-knockout (*Stim1-*smKO) mice. VSMCs from *Stim1-*smKO mice also exhibited a reduction in nanoscale colocalization between Ca^2+^-release sites on the SR and Ca^2+^-activated ion channels on the PM, accompanied by diminished channel activity. *Stim1-*smKO mice were hypotensive and resistance arteries isolated from them displayed blunted contractility. These data suggest that STIM1 – independent of SR Ca^2+^ store depletion – is critically important for stable peripheral coupling in contractile VSMCs.

## Introduction

Subcellular Ca^2+^-signaling microdomains formed by interactions between the sarcoplasmic reticulum (SR) and the plasma membrane (PM) are vital for many physiological processes, including regulation of the contractility of vascular smooth muscle cells (VSMCs) (1, 2). Ca^2+^ signals that occupy these compartments are typified by Ca^2+^ sparks – large-amplitude Ca^2+^ transients that reflect optically detected Ca^2+^ ions released into the cytosol from the SR through clusters of type 2 ryanodine receptors (RyR2s). Ca^2+^ sparks activate clusters of large-conductance Ca^2+^-activated K^+^ (BK) channels on the PM, generating transient, macroscopic outward K^+^ currents that hyperpolarize the PM (1, 3, 4). A complementary Ca^2+^ signaling pathway that causes VSMC membrane depolarization and elevated contractility is formed by interactions between inositol 1,4,5-trisphosphate receptors (IP3Rs) on the SR and monovalent cation-selective, Ca^2+^-activated TRPM4 (transient receptor potential melastatin 4) channels on the PM. Ca^2+^ released from the SR through IP3Rs activates Na^+^ influx through TRPM4, causing depolarization of the PM and increased VSMC contractility (2, 5). The close association of the SR and PM creates subcellular compartments where the local Ca^2+^ ion concentration can reach the micromolar range required for activation of BK and TRPM4 channels under physiological conditions (6). In non-excitable cells, endoplasmic reticulum (ER)-PM junctions and associated proteins have been well characterized (7, 8). In contrast, SR-PM junctional areas of VSMCs and the essential proteins that mediate these interactions remain poorly understood.

The ER-PM junctions of non-excitable cells are highly specialized hubs for ion channel signaling cascades. These spaces are the sites of one of the most ubiquitous receptor-regulated Ca^2+^ entry pathway in such cells, termed store-operated Ca^2+^ entry (SOCE), which is mediated by the ER-resident Ca^2+^-sensing protein STIM1 (stromal interaction molecule 1) and Ca^2+^-selective channels of the Orai group on the PM (9–11). STIM1 is a single-pass transmembrane ER/SR protein that possesses a low-affinity Ca^2+^-sensing EF-hand facing the lumen of the ER/SR (9, 12–30). Following store depletion by IP3-producing receptor agonists, STIM1 acquires an extended conformation and migrates to ER-PM junctions, exposing a cytosolic STIM-Orai activating region that physically traps and activates Orai channels on the PM (9, 10, 12, 18, 22, 24, 26, 29–35). The other STIM protein family member, STIM2, is structurally similar to STIM1. Fully differentiated VSMCs from systemic arteries express STIM1 but not STIM2 and do not exhibit detectable SOCE (36–38). Many species express IP3Rs but lack STIM and Orai proteins, suggesting that receptor-evoked Ca^2+^ signaling is not always complemented by the operation of STIM and Orai mechanisms (39). Evolutionary evidence indicates that Orai appeared before STIM, implying that STIM might have arisen to support the function of ER-PM junctions and only subsequently co-opted an existing Orai for SOCE (39). Additional accumulating evidence indicates that, in addition to its role in SOCE, mammalian STIM1 protein serves as an essential regulator of several other ion channels and signaling pathways. STIM1 both positively and negatively regulates the function of L-type voltage-gated Ca^2+^ channels (Cav1.2) (40), transient receptor potential canonical (TRPC) channels (41), and arachidonate-regulated Ca^2+^ (ARC) channels (42). It has also been reported to regulate the function of Ca^2+^ pumps, such as the SR/ER Ca^2+^ ATPase (SERCA) and PM Ca^2+^ ATPase (PMCA) as well as several cAMP-producing adenylyl cyclases at the PM (43–46).

In the current study, we investigated the role of STIM1 in the formation of stable peripheral coupling sites in native, contractile SMCs from cerebral arteries. We show that STIM1 is required for stable interactions between the SR and PM. We further show that this function of STIM1 is independent of SR Ca^2+^ store depletion and acts to sustain subcellular Ca^2+^ signaling pathways that are essential for the regulation of VSMC contractility.

## Results

### Stim1-smKO mice lack STIM1 protein expression in VSMCs

Mice with *loxP* sites flanking exon 2 of the *Stim1* gene (*Stim1^fl/f^*^l^ mice) were crossed with myosin heavy chain 11 (*Myh11*)-*cre*/ERT2 mice (47, 48), generating *Myh11-Cre-Stim1^fl/wt^* mice, in which *Myh11* promoter-driven *cre* expression is induced by injection of tamoxifen. Heterozygous *Myh11-Cre-Stim1^fl/wt^* mice were then intercrossed, yielding tamoxifen-inducible, SMC-specific *Stim1-*knockout mice (*Myh11*-*Cre*-*Stim1^fl/fl^*), hereafter termed *Stim1-*smKO mice. Cre-recombinase expression was induced in male *Stim1-*smKO mice by daily intraperitoneal injection of tamoxifen (100 µL, 10 mg/mL) for 5 days, beginning at 4–6 weeks of age. Controls for all experiments consisted of *Stim1-*smKO mice injected with the vehicle for tamoxifen (sunflower oil). Mice were used for experiments 1 week after the final injection. The Wes capillary electrophoresis immunoassay-based protein detection system was used for qualitative and quantitative assessment of STIM1 protein in smooth muscle tissues from *Stim1-*smKO and control mice. STIM1 protein was readily detected as a single band in cerebral artery, mesenteric artery, aortic, colonic, and bladder smooth muscle isolated from control mice, but was virtually undetectable in smooth muscle isolated from tamoxifen-injected *Stim1-*smKO mice (Figure 1A). STIM1 protein levels normalized to total protein (Supplementary figure 1A) were significantly lower in cerebral artery, aortic, colonic, and bladder smooth muscle from tamoxifen-injected *Stim1-*smKO mice compared with controls (Figure 1A). In contrast, STIM1 protein expression was detected at similar levels in whole brains from both control and tamoxifen-injected *Stim1-*smKO mice (Figure 1A), reflecting STIM1 expression in brain cells apart from VSMCs. Tamoxifen injection had no effect on STIM1 protein levels in *Myh-11-Cre*-positive *Stim1^wt/wt^* mice (Supplementary figure 1B–G).

**Figure 1:**
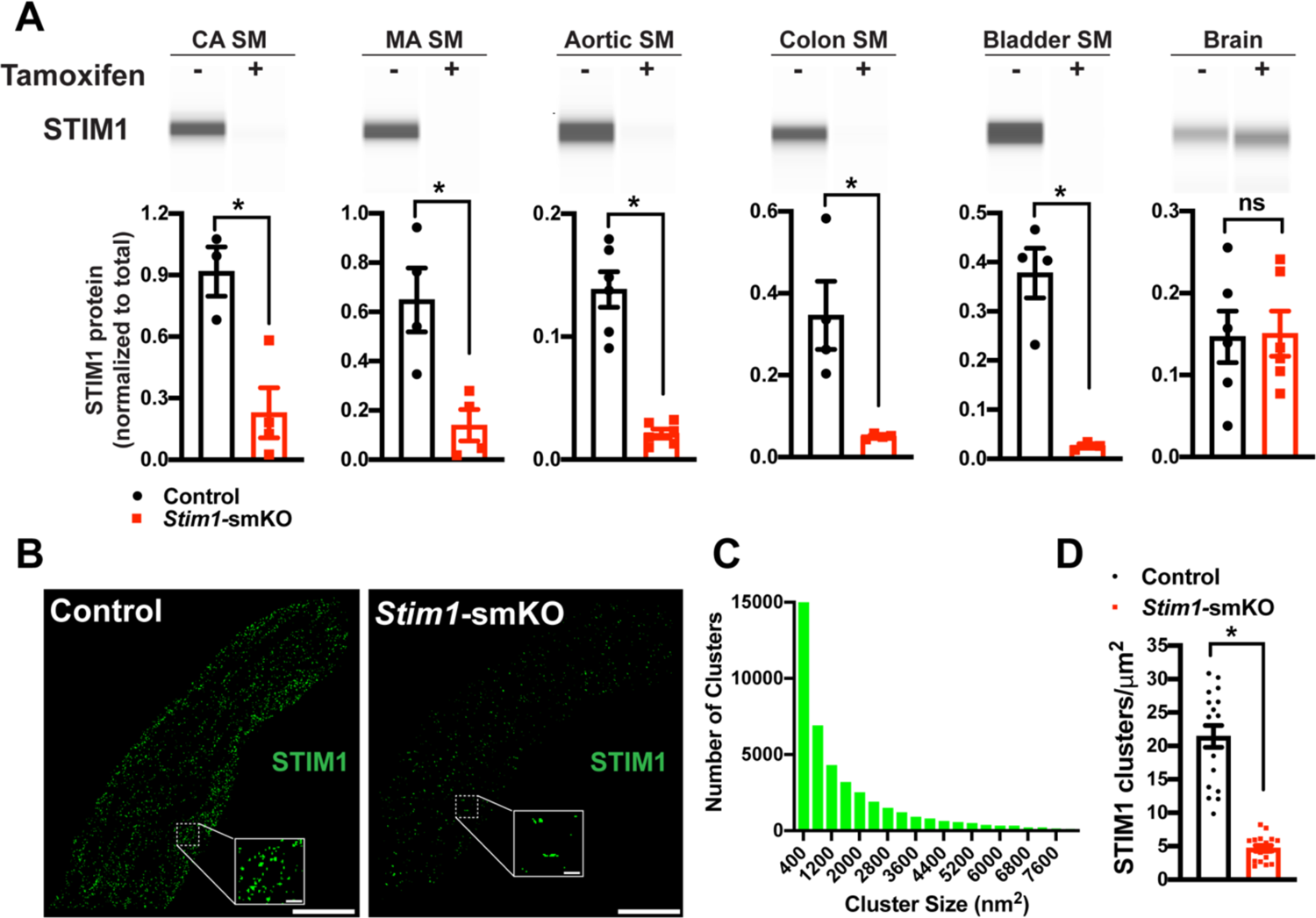
Inducible, SMC-specific *Stim1* knockout. **(A)** Representative Wes protein capillary electrophoresis experiments, presented as Western blots, showing STIM1 protein expression levels in smooth muscle tissues and brains of vehicle- and tamoxifen-injected *Stim1-*smKO mice. Summary data showing densitometric analyses of STIM1 protein expression in cerebral artery smooth muscle (CA SM), mesenteric artery smooth muscle (MA SM), aortic smooth muscle, colonic smooth muscle, bladder smooth muscle and brain, normalized to total protein (n = 3–6 mice/group; *P < 0.05, unpaired t-test). ns, not significant. **(B)** Representative superresolution localization maps of isolated cerebral artery SMCs from control and *Stim1-*smKO mice immunolabeled for STIM1. Insets: enlarged areas highlighted by the white squares in the main panels. Scale bars: 3 µm (main panels) and 250 nm (inset panels). **(C)** Distribution plot of the surface areas of individual STIM1 clusters in cerebral artery SMCs isolated from control mice (n = 42726 clusters from 18 cells from 3 mice). **(D)** STIM1 cluster density in cerebral artery SMCs isolated from control and *Stim1-*smKO mice (n = 18 cells from 3 mice/group; *P < 0.05, unpaired t-test).

In further studies, single SMCs from cerebral arteries isolated from control and *Stim1-*smKO mice were enzymatically dispersed, immunolabeled with an anti-STIM1 primary antibody, and imaged using a GSDIM (ground state depletion followed by individual molecule return) superresolution microscopy system, which we previously showed using DNA-origami-based nanorulers has a lateral resolution of 20–40 nm (49–51). VSMCs from control mice exhibited punctate STIM1 protein clusters (Figure 1B). Frequency analyses revealed that the sizes of these clusters were exponentially distributed, with a majority of clusters (∼95%) ranging in area between 400 and 7600 nm^2^ (mean = 2135 ± 21 nm^2^; median = 800 nm^2^) (Figure 1C). STIM1 cluster density was significantly reduced in VSMCs isolated from *Stim1-*smKO mice (Figure 1D). The number of GSDIM events in VSMCs isolated from tamoxifen-injected *Stim1-*smKO mice was comparable to background levels observed in cells from control mice immunolabeled with secondary antibody only, providing further evidence of effective STIM1 knockdown (Supplementary figure 2A and B). Taken together, these data demonstrate selective, tamoxifen-inducible SMC-specific knockout of STIM1 expression in *Stim1-*smKO mice.

**Figure 2:**
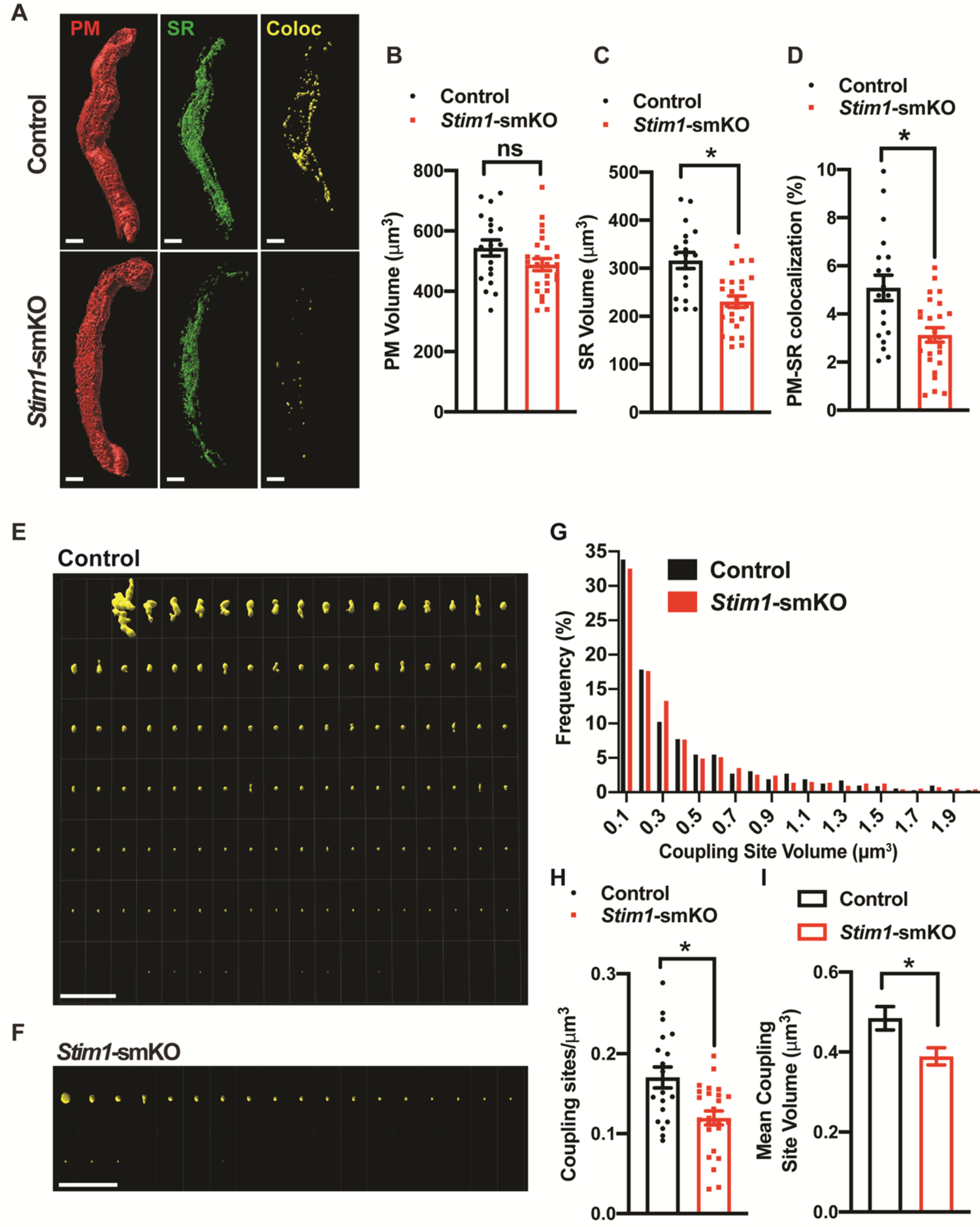
*Stim1* knockout decreases the density and area of PM-SR coupling sites. **(A)** Representative 3D surface reconstructions of cerebral artery SMCs isolated from control *Stim1-*smKO mice labeled with PM (red) and SR (green) dyes and imaged using SIM. Representations of colocalizing PM and SR surfaces (yellow), generated from surface reconstructions. Scale bar: 5 µm. **(B** and **C)** PM and SR volumes and **(D)** PM-SR colocalization (%) in cells from control and *Stim1*-smKO mice. **(E** and **F)** Ensemble images of all PM-SR colocalization sites in single cells from the control and *Stim1*-smKO mice shown in panel A. Scale bar: 10 µm **(G)** Frequency distribution of the volumes of individual PM-SR colocalization sites in VSMCs isolated from control and *Stim1-*smKO mice. **(H)** Densities and **(I)** mean volumes of individual coupling sites in VSMCs from control and *Stim1-*smKO mice. Data are for 1736 colocalization sites in 19 cells from 6 mice for control and 1484 colocalization sites in 25 cells from 7 mice for *Stim1*-smKO (*P < 0.05, unpaired t-test). ns, not significant.

### PM and SR coupling is diminished in VSMCs from Stim1-smKO mice

To investigate how STIM1 knockout affects PM and SR interactions, we costained native SMCs isolated from cerebral arteries of control and *Stim1-*smKO mice with Cell-Mask Deep Red and ER-Tracker green to label the PM and SR, respectively, as described in our prior publications (49, 52). Using live-cell structured illumination microscopy (SIM), we acquired Z-stack images of PM- and SR-labeled VSMCs as 0.25-µm slices. We then reconstructed the 3D surfaces of the PM and SR from these images (Figure 2A), also generating a third surface indicating the sites of colocalization between the PM and SR (Figure 2A; Supplementary movies 1 and 2). The mean volume of the PM did not differ between *Stim1-*smKO and control mice, but the volume of the SR was smaller in cells isolated from *Stim1-*smKO mice (Figure 2B and C). The overall PM-SR colocalization was significantly reduced in VSMCs from *Stim1-*smKO mice compared with controls (Figure 2D). As shown in representative image galleries of individual colocalization sites (Figure 2E), the majority of PM-SR coupling sites in cells from both groups formed spherical surfaces, but some of the larger structures exhibited an elongated morphology. Frequency analyses showed that the volume of individual colocalization sites in cells from both groups exhibited an exponential distribution (Figure 2G). In addition, the number of coupling sites per unit volume and mean volume of individual sites were smaller in cells from *Stim1-*smKO mice compared with those from controls (Figure 2H and I). These data indicate that interactions between the PM and SR are decreased by *Stim1* knockout in VSMCs.

### Stim1 knockout decreases colocalization of BK and RyR2 protein clusters

BK channels on the PM of VSMCs are functionally coupled with RyR2s on the SR (1). Therefore, we investigated how *Stim1* knockout affects the nanoscale structure of the BK-RyR2 signaling complex using GSDIM superresolution microscopy. Freshly isolated VSMCs from *Stim1-*smKO and control mice were co-immunolabeled for RyR2 and the BK channel pore-forming subunit BK*α* and imaged using GSDIM in epifluorescence illumination mode. The resulting superresolution localization maps (Figure 3A, left-most panels) showed that both proteins were present as defined clusters in VSMCs. Using an objects-based analysis (OBA) approach (53, 54) as described in previous publications (49, 51, 52, 55, 56), we generated new maps of RyR2 clusters that overlapped at the resolution limit of our microscope system (∼20-40 nm) with the centroid of each BK cluster, and BK clusters that overlapped with the centroid of each RyR2 cluster. These two maps were then merged to reveal colocalized RyR2-BK channel protein clusters in VSMCs from both groups of animals that were below the resolution of our GSDIM system (Figure 3A, middle and right-most panels). Particle analysis of these clusters showed that the density of individual BK protein clusters (number of clusters per unit area) was similar for both groups of animals (Figure 3B), whereas the density of individual RyR2 clusters was lower in VSMCs from *Stim1-*smKO mice compared with controls (Figure 3C). In both groups, the sizes of individual BK channel and RyR2 clusters followed an exponential distribution (Figure 3B and C). The mean size of individual BK clusters was smaller in VSMCs from *Stim1-*smKO mice compared with those from controls (Figure 3B); in contrast, the mean size of RyR2 clusters was slightly larger in cells from *Stim1-*smKO mice (Figure 3C). In terms of colocalization, this analysis showed a significant reduction in the density of colocalized BK-RyR2 protein clusters in VSMCs from *Stim1-*smKO mice compared with controls (Figure 3D). The mean size of colocalizing clusters from *Stim1-*smKO mice was smaller compared with those from control mice (Figure 3D), and the sizes of BK-RyR2 colocalization sites in cerebral artery SMCs from both groups exhibited an exponential distribution (Figure 3D). These data indicate that *Stim1* knockout decreases the frequency and size of close contact sites between RyR2 and BK channel protein clusters in VSMCs.

**Figure 3:**
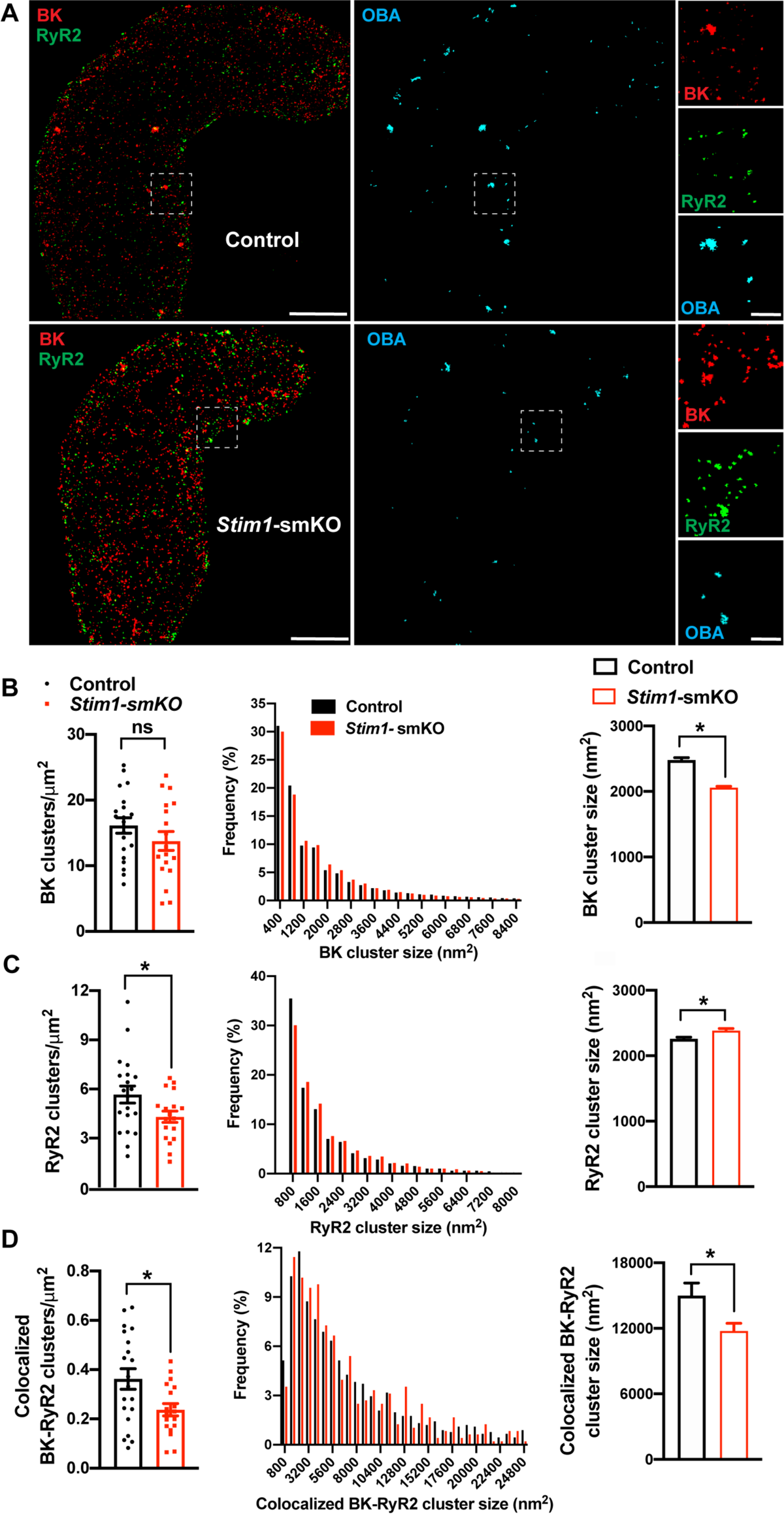
*Stim1* knockout decreases colocalization of BK and RyR2 protein clusters. **(A)** Superresolution localization maps of freshly isolated cerebral artery SMCs from control and *Stim1-*smKO mice immunolabeled for BK (red) and RyR2 (green). Colocalized BK and RyR2 clusters were identified by object-based analysis (OBA) and mapped (cyan). Scale bar: 3 µm. Panels to the right show enlarged areas of the original superresolution maps indicated by the white boxes. Scale bar: 500 nm. **(B)** Summary data showing the density (clusters per unit area), frequency distribution of sizes, and mean size of BK channel clusters. **(C)** Summary data showing the density, frequency distribution of sizes, and mean size of RyR2 clusters. **(D)** Summary data showing the density, frequency distribution of sizes, and mean size of colocalizing BK and RyR2 clusters, identified using object-based analysis. For density data, n = 20 cells from 3 mice for controls and n = 18 cells from 3 mice for *Stim1-*smKO mice. For frequency distribution and mean cluster size data: control, n = 44,340 BK channel clusters, n = 15,193 RyR2 clusters, and n = 1054 colocalizing clusters; Stim1-smKO: n = 30,552 BK channel clusters, n = 9702 RyR2 clusters, and n = 547 colocalizing clusters (*P < 0.05, unpaired t-test). ns, not significant.

### Stim1 knockout decreases colocalization of TRPM4 and IP3R protein clusters

TRPM4 channels on the PM are functionally coupled with IP3Rs on the SR (2). Therefore, we also investigated how interactions between PM TRPM4 channels and SR IP3Rs were altered by *Stim1* knockout. Freshly isolated VSMCs from control and *Stim1-*smKO mice were co-immunolabeled for TRPM4 and IP3R and imaged using GSDIM in epifluorescence illumination mode. The resulting GSDIM localization maps showed that these proteins are present as discrete clusters in cells (Figure 4A, left-most panels).

**Figure 4:**
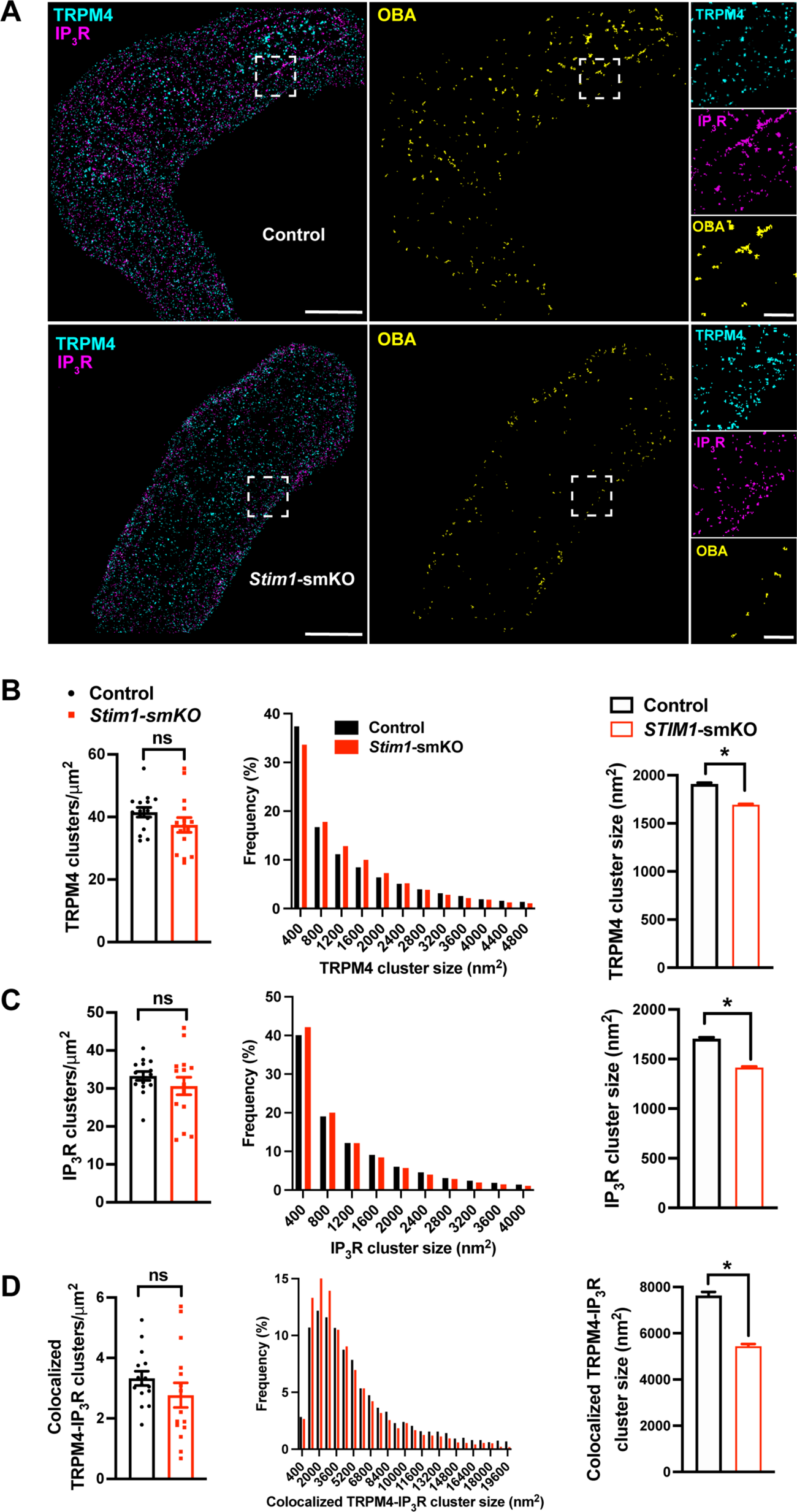
*Stim1* knockout decreases colocalization of TRPM4 and IP3R protein clusters. **(A)** Superresolution localization maps of freshly isolated cerebral artery SMCs from control and *Stim1-*smKO mice immunolabeled for TRPM4 (cyan) and IP3R (magenta). Colocalized TRPM4 and IP3R clusters were identified by object-based analysis (OBA) and mapped (yellow). Scale bar: 3 µm. Panels to the right show enlarged areas of the original superresolution maps indicated by white boxes. Scale bar: 500 nm. **(B)** Summary data showing the density (clusters per unit area), frequency distribution of sizes, and mean size of TRPM4 channel protein clusters. **(C)** Summary data showing the density, frequency distribution of sizes, and mean size of IP3R clusters. **(D)** Summary data showing the density, frequency distribution of sizes, and mean size of colocalizing TRPM4 and IP3R clusters, identified using object-based analysis. For density data, n = 15 cells from 3 mice for both control and *Stim1-*smKO mice. For frequency distribution and mean cluster size data: control, n = 64292 TRPM4 channel clusters, n = 51728 IP3R clusters, and n = 5164 colocalizing clusters; *Stim1-*smKO mice, n = 56771 TRPM4 channel clusters, n = 45717 IP3R, and n = 3981 colocalizing clusters (*P<0.05, unpaired t-test). ns, not significant.

We next used OBA to identify and map individual and colocalized TRPM4 and IP3R protein clusters (Figure 4A, middle and right-most panels). This analysis showed that the densities of individual TRPM4 and IP3R clusters were similar in both groups (Figure 4B and C) and that their sizes were exponentially distributed (Figure 4B and C). The mean sizes of individual TRPM4 and IP3R clusters were smaller in VSMCs from *Stim1-*smKO mice compared with those from controls. (Figure 4B and C). The density of colocalized TRPM4-IP3R cluster sites did not differ between groups (Figure 4D, left), but the sizes of these colocalized clusters were smaller in cells from *Stim1-*smKO mice compared with those from controls (Figure 4D, middle). Like individual clusters, colocalized clusters exhibited an exponential distribution (Figure 4D, right).

### Stim1 knockout alters the properties of Ca^2+^ sparks

To investigate how *Stim1* knockout alters fundamental Ca^2+^ signaling mechanisms, we loaded freshly isolated VSMCs with the Ca^2+^-sensitive fluorophore Fluo-4-AM and imaged them using live-cell, high-speed, high-resolution spinning-disk confocal microscopy. Spontaneous Ca^2+^ sparks were present in cerebral artery SMCs from both control (Figure 5A; Supplementary movie 3) and *Stim1-*smKO (Figure 5B; Supplementary movie 4) mice. The frequency of Ca^2+^ spark events did not differ between groups (Figure 5C). However, the mean amplitude of Ca^2+^ spark events was significantly greater in VSMCs isolated from *Stim1-*smKO mice compared with those from controls (Figure 5D). Further analyses revealed that spatial spreads, durations, and decay times of individual Ca^2+^ spark events were significantly greater in VSMCs isolated from *Stim1-*smKO mice compared with those taken from control mice, but rise times did not differ (Figure 5E–H). To investigate the effects of *Stim1* knockout on total SR Ca^2+^ store load, we applied a bolus of caffeine (10 mM) to Fluo-4-AM–loaded VSMCs isolated from control and *Stim1-*smKO mice. The peak amplitude of caffeine-evoked global increases in cytosolic [Ca^2+^] did not differ between groups (Figure 5I), indicating that *Stim1* knockout did not alter total SR [Ca^2+^]. Therefore, alterations in the properties of Ca^2+^ sparks associated with the knockout of *Stim1* are not the result of changes in SR Ca^2+^ load.

**Figure 5:**
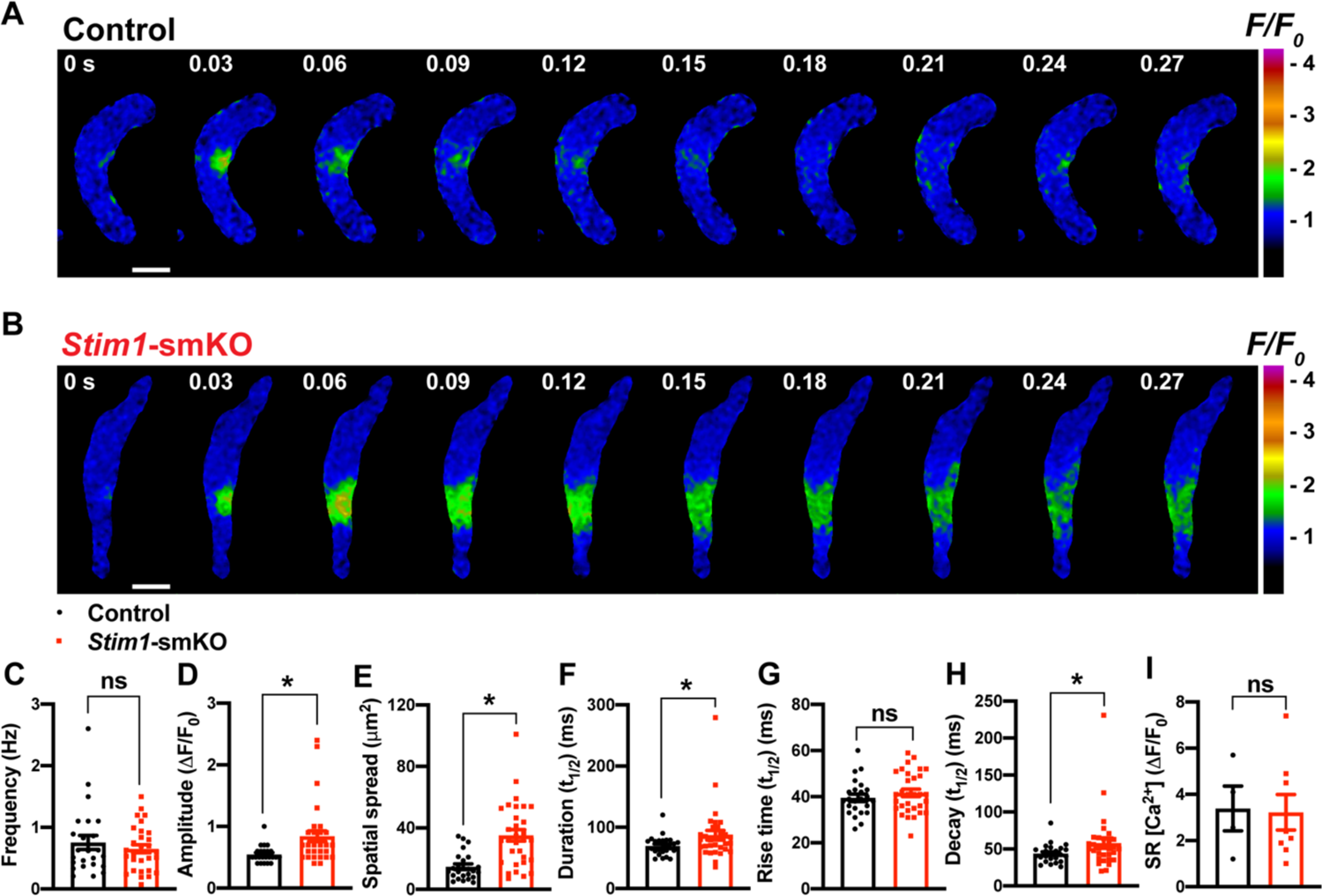
*Stim1* knockout alters Ca^2+^ spark properties. **(A** and **B)** Representative time-course images of cerebral artery SMCs isolated from a control **(A)** or *Stim1-*smKO **(B)** mouse exhibiting Ca^2+^ spark events, presented as changes in fractional fluorescence (F/F0). The elapsed time of the event is shown in seconds (s). Scale bar: 10 µm. **(C–H)** Summary data showing Ca^2+^ spark frequency **(C)**, amplitude **(D)**, spatial spread **(E)**, event duration **(F)**, rise time **(G)**, and decay time **(H)** in VSMCs isolated from control and *Stim1-*smKO mice (control, n = 24 spark sites in 12 cells from 2 mice; *Stim1-*smKO, n = 31 spark sites in 12 cells from 2 mice; *P < 0.05, unpaired t-test). ns, not significant. **(I)** Summary data showing caffeine (10 mM)-evoked changes in global Ca^2+^ in cerebral artery SMCs isolated from control and *Stim1-*smKO mice. (control, n = 4 cells from 2 mice; *Stim1-*smKO, n = 8 cells from 2 mice, unpaired t-test). ns, not significant.

### Stim1 knockout diminishes physiological BK and TRPM4 channel activity

We next used patch-clamp electrophysiology to investigate how knockout of *Stim1* affects the activity of BK and TRPM4 channels in VSMCs. When Ca^2+^ sparks activate clusters of BK channels at the PM, they generate macroscopic K^+^ currents termed spontaneous transient outward currents (STOCs) (1). Here, we recorded STOCs over a range of membrane potentials using the amphotericin B perforated patch-clamp configuration, which allows the membrane potential to be controlled without disrupting intracellular Ca^2+^ signaling pathways (49, 52). The frequencies and amplitudes of STOCs were lower in VSMCs from *Stim1-*smKO mice compared with those from controls at all membrane potentials greater than −60 mV (Figure 6A, B and C). To determine if diminished STOC activity was attributable to a decrease in the total number of BK channels available for activation at the PM, we measured whole-cell BK channel currents. Cerebral artery SMCs isolated from *Stim1-*smKO and control mice were patch-clamped in the conventional whole-cell configuration, and whole-cell K^+^ currents were recorded during application of voltage ramps. Using the selective BK blocker paxilline to isolate BK channel currents, we found that whole-cell BK current amplitude did not differ between VSMCs from control and *Stim1-*smKO mice (Figure 6D and E), indicating that the number of BK channels available for activation and their functionality was not altered by *Stim1* knockout. Also, *Stim1* knockout did not alter mRNA levels of BK *α*- or *β*1-subunits or RyR2s in cerebral arteries (Supplementary figure 3A). These findings indicate that diminished STOC activity following knockout of *Stim1* may result from impaired functional coupling of RyR2 with BK channels.

**Figure 6:**
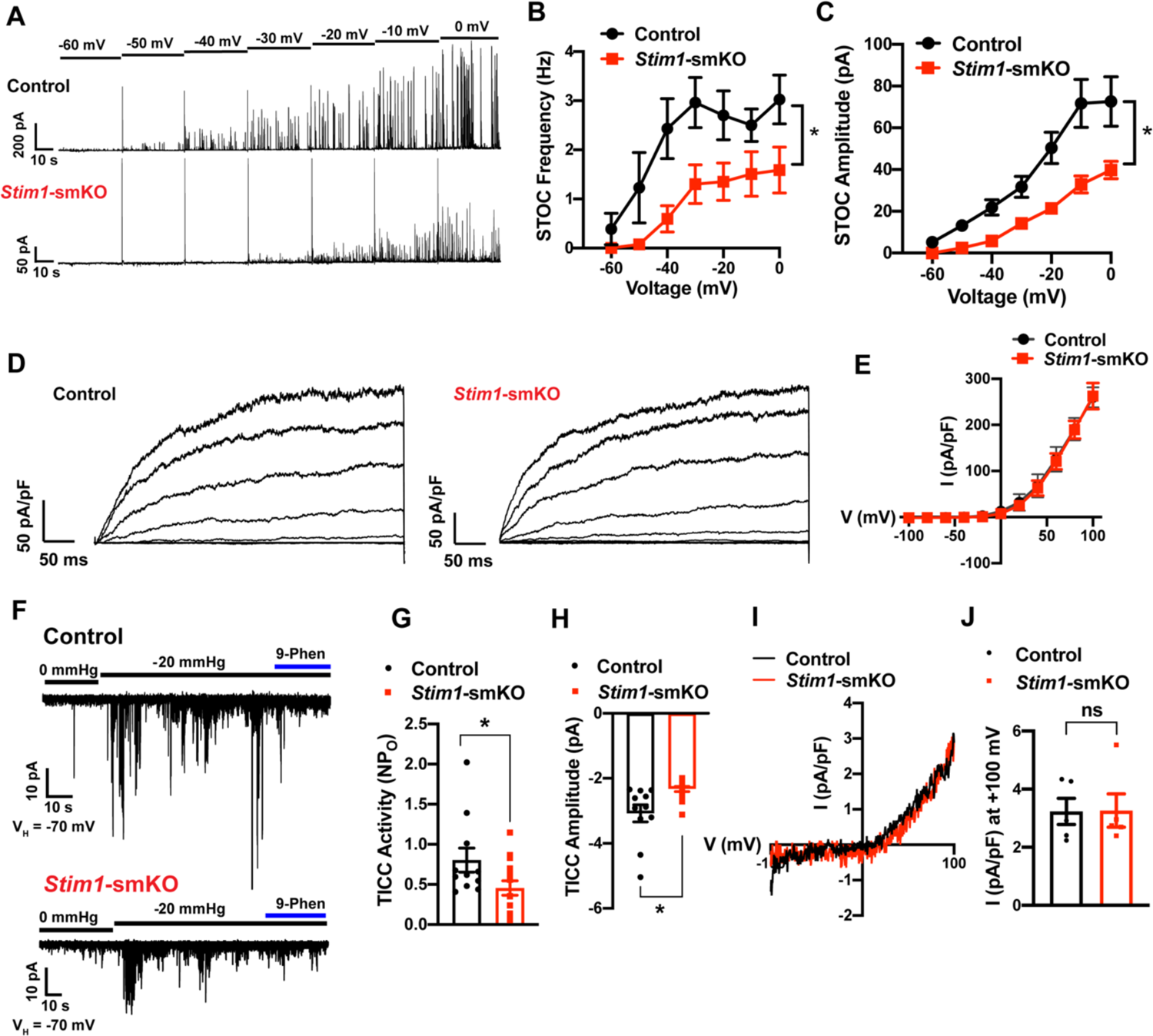
*Stim1* knockout diminishes physiological BK and TRPM4 channel activity. **(A)** Representative traces of STOCs in cerebral artery SMCs from control and *Stim1-*smKO mice, recorded by patch-clamping in the perforated-patch configuration over a range of membrane potentials (−60 to 0 mV). **(B** and **C)** Summary data showing STOC frequency **(B)** and amplitude **(C)** (control, n = 13 cells from 4 animals; *Stim1-*smKO, n = 17 cells from 5 mice; *P < 0.05, two-way ANOVA). **(D)** Representative traces of paxilline (1 μM)-sensitive BK currents in cerebral artery SMCs from control and *Stim1-*smKO mice, recorded by patch-clamping in conventional whole-cell mode during a series of command voltage steps (−100 to +100 mV). **(E)** Summary data for whole-cell BK currents (control, n = 6 cells from 3 mice; *Stim1-*smKO, n = 7 cells from 3 mice; two-way ANOVA). **(F)** Representative traces of TRPM4 currents in cerebral artery SMCs from control and *Stim1-*smKO mice voltage-clamped at −70 mV, recorded by patch-clamping in the perforated-patch configuration. TRPM4 currents were evoked as TICCs by application of negative pressure (−20 mmHg) through the patch pipette and were blocked by bath-application of 9-phenanthrol (9-phen; 30 μM). **(G)** Summary data showing TICC activity as TRPM4 channel open probability (*NP*o) and **(H)** TICC amplitude in control (*n* = 12 cells from 5 mice) and *Stim1-*smKO (*n* = 15 cells from 5 mice) mice (*P < 0.05, unpaired t-test). **(I)** Representative conventional whole-cell patch-clamp recordings of 9-phenanthrol–sensitive TRPM4 currents in cerebral artery SMCs from control and *Stim1-*smKO mice. Currents were activated by free Ca^2+^ (200 µM), included in the patch pipette solution, and were recorded using a ramp protocol from −100 to 100 mV from a holding potential of −60 mV. **(J)** Summary of whole-cell TRPM4 current density at +100 mV (control, n = 5 cells from 3 mice; *Stim1-*smKO, n = 5 cells from 3 mice, unpaired t-test). ns, not significant.

TRPM4 is a Ca^2+^-activated, monovalent cation-selective channel that is impermeable to divalent cations (57). At membrane potentials in the physiological range for VSMCs (−70 to −30 mV), TRPM4 channels conduct inward Na^+^ currents that depolarize the plasma membrane in response to increases in intraluminal pressure and receptor-dependent vasoconstrictor agonists (58, 59). Under native conditions, TRPM4 channels are activated by Ca^2+^ released from the SR through IP3Rs, generating transient inward cation currents (TICCs) (2, 60). To determine the effects of STIM1 knockout on TRPM4 activity, we recorded TICCs using the amphotericin B perforated patch-clamp configuration (60). In agreement with previous reports (5, 59), we found that TICC activity in VSMCs from control mice was increased following application of negative pressure (−20 mmHg) through the patch pipette to stretch the plasma membrane, an effect that was attenuated by the selective TRPM4 blocker, 9-phenanthrol (Figure 6F). TICC activity and amplitude in VSMCs isolated from *Stim1-*smKO mice were significantly reduced compared with those from controls (Figure 6F, G, and H). To determine if these differences were attributable to changes in TRPM4 channel function or availability, we activated TRPM4 currents in VSMCs from *Stim1-*smKO and control mice using an internal solution containing 200 µM free Ca^2+^ and compared whole-cell TRPM4 currents in both groups by patch-clamping VSMCs in the conventional whole-cell configuration (61). The TRPM4-sensitive component of the current was isolated by applying 9-phenanthrol. We found that whole-cell TRPM4 current amplitudes did not differ between VSMCs from control and *Stim1-*smKO mice (Figure 6I and J), suggesting that the number of TRPM4 channels available for activation at the PM was not altered by *Stim1* knockout. In addition, *Stim1* knockout did not alter mRNA levels of TRPM4 subunits or any of the IP3R subtypes (1,2, or 3) in cerebral arteries (Supplementary figure 3B). These findings suggest that diminished TICC activity following knockout of *Stim1* results from impaired functional coupling of IP3Rs with TRPM4 channels.

### The contractility of resistance arteries from Stim1-smKO mice is blunted

Knockout of *Stim1* in VSMCs decreased the activity of BK and TRPM4 channels under physiological recording conditions. These channels have opposing effects on VSMC membrane potential, contractility and arterial diameter, with BK channels causing dilation (1) and TRPM4 channels causing constriction (62). Thus, the overall functional impact of deficient channel activity is not immediately apparent. Therefore, to investigate the net consequences of *Stim1* knockout on arterial contractile function, we employed a series of *ex vivo* pressure myography experiments. Constrictions of intact cerebral pial arteries in response to a depolarizing concentration (60 mM) of extracellular KCl did not differ between groups (Figure 7A), suggesting that knocking out *Stim1* in cerebral artery SMCs did not grossly alter voltage-dependent Ca^2+^ influx or underlying contractile processes. Contractile responses to increases in intraluminal pressure (myogenic vasoconstriction) were evaluated by measuring steady-state luminal diameter at intraluminal pressures over a range of 5 to 140 mmHg in the presence (active response) and absence (passive response) of extracellular Ca^2+^. Myogenic tone, calculated by normalizing active constriction to passive dilation, was significantly lower in cerebral arteries from *Stim1-*smKO mice compared with those from controls (Figure 7B and C). Contractile responses to the synthetic thromboxane A2 receptor agonist U46619 were also significantly blunted in cerebral arteries from *Stim1-*smKO mice compared with those from vehicle-treated controls (Figure 7D and E). These data demonstrate that the ability of cerebral arteries from *Stim1-*smKO mice to contract in response to physiological stimuli is impaired. Additional investigations using 3^rd^-order mesenteric arteries yielded similar findings (Figure 7F–J), indicating widespread vascular dysfunction in *Stim1-*smKO mice.

**Figure 7:**
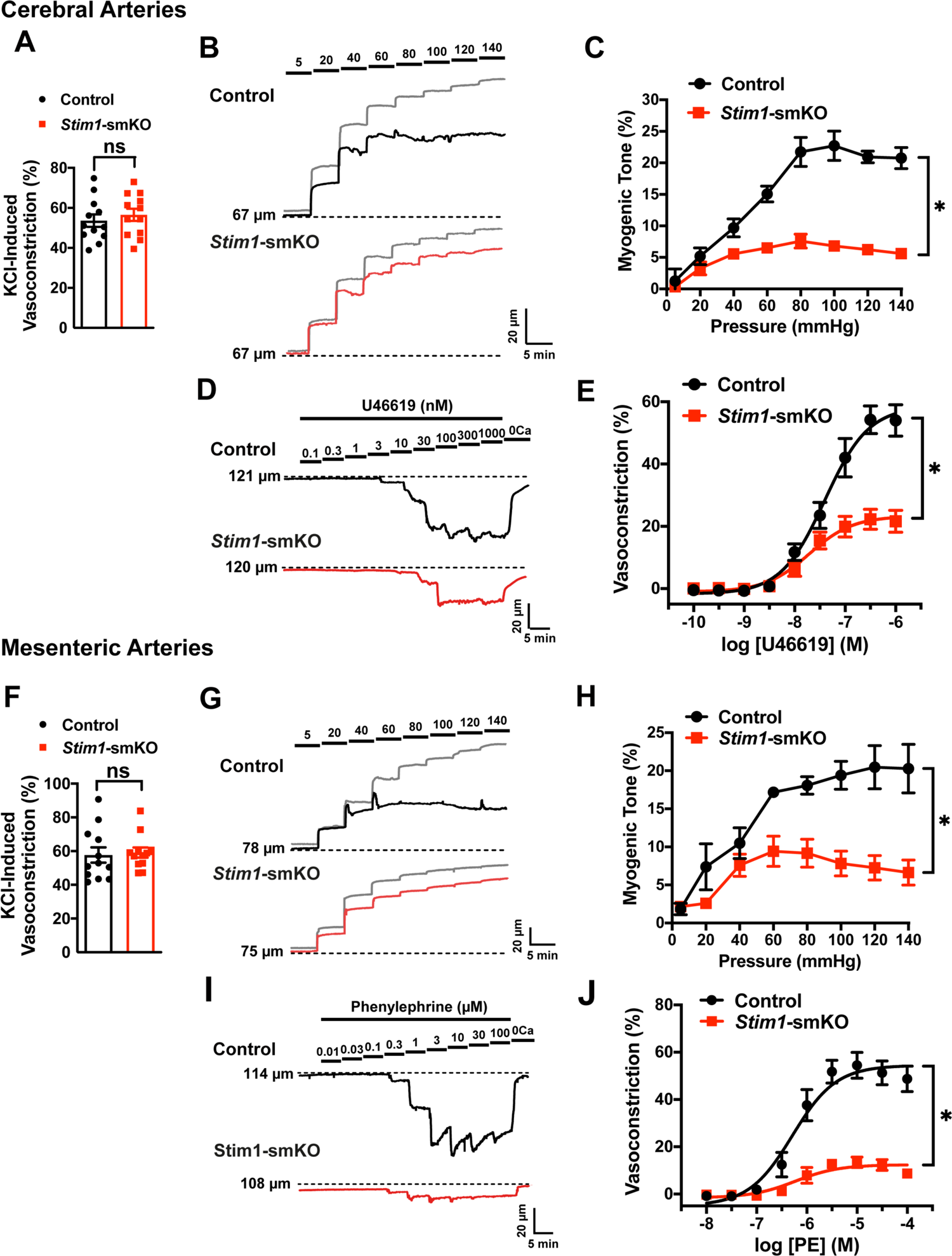
Resistance arteries from *Stim1-*smKO mice are dysfunctional. **(A)** Summary data showing vasoconstrictor responses to 60 mM KCl in cerebral pial arteries isolated from control and *Stim1-*smKO mice (n = 12 vessels from 6 mice for both groups, unpaired t-test). ns, not significant. **(B)** Representative traces showing changes in luminal diameter over a range of intraluminal pressures (5 to 140 mmHg) in cerebral pial arteries isolated from control (black trace) and *Stim1-*smKO (red) mice. Gray traces represent passive responses to changes in intraluminal pressure for each artery. **(C)** Summary data showing myogenic reactivity as a function of intraluminal pressure (n = 6 vessels from 3 mice for each group; *P < 0.05, two-way ANOVA). **(D)** Representative traces showing changes in luminal diameter in response to a range of concentrations (0.1 to 1000 nM) of the vasoconstrictor agonist U46619 in cerebral arteries isolated from control (black trace) and *Stim1-*smKO (red trace) mice. (**E**) Summary data showing vasoconstriction as a function of U46619 concentration (n = 6 vessels from 3 mice for each group; *P < 0.05, two-way ANOVA). **(F)** Summary data showing vasoconstrictor responses to 60 mM KCl in 3^rd^-order mesenteric arteries isolated from control and *Stim1-*smKO mice (n = 12 vessels from 6 mice for both groups, unpaired t-test). ns, not significant **(G)** Representative traces showing changes in luminal diameter over a range of intraluminal pressures (5 to 140 mmHg) in 3^rd^-order mesenteric arteries isolated from control (black trace) and *Stim1-*smKO (red) mice. Gray traces represent passive responses to changes in intraluminal pressure for each artery. **(H)** Summary data for myogenic reactivity as a function of intraluminal pressure (n = 6 vessels from 3 mice for each group, 2-way ANOVA, *P < 0.05). (**I**) Representative traces showing changes in luminal diameter in response to a range of concentrations (0.01 to 100 μM) of the vasoconstrictor agonist phenylephrine (PE) in 3^rd^-order mesenteric arteries isolated from control (black trace) and *Stim1-*smKO (red trace) mice. **(J)** Summary data for vasoconstriction as a function of PE concentration, presented as means ± SEM (n = 6 vessels from 3 mice for each group; *P < 0.05, two-way ANOVA).

### Stim1-smKO mice are hypotensive

Age-matched *Stim1*-smKO mice were surgically implanted with radio telemetry transmitters as previously described (63). After a recovery period (14 days), systolic and diastolic blood pressure (BP), heart rate (HR), and locomotor activity levels were recorded for 48 hours before tamoxifen injection (control). Systolic and diastolic BP, HR, and activity levels were again recorded for 48 hours, beginning 1 week after completing the tamoxifen injection protocol (*Stim1*-smKO). Normal diurnal variations were observed for all parameters (Figure 8). The mean systolic BP of *Stim1*-smKO mice was lower than that of control mice during both day and night cycles (Figure 8A), whereas diastolic BP did not differ between groups (Figure 8B). Mean arterial pressure (MAP) (Figure 8C) was lower in *Stim1*-smKO mice compared with controls at night, and trends to be different during the day (P = 0.056). The pulse pressure of *Stim1*-smKO mice was lower than that of control mice during both day and night cycles (Figure 8D). HR (Figure 8E) and locomotor activity (Figure 8F) did not differ between groups. Injection of vehicle did not affect BP, HR, or locomotor activity (Supplementary Figure 4). These data indicate that acute knockout of *Stim1* in SMCs lowers BP, probably due to diminished arterial contractility and decreased total peripheral resistance.

**Figure 8.**
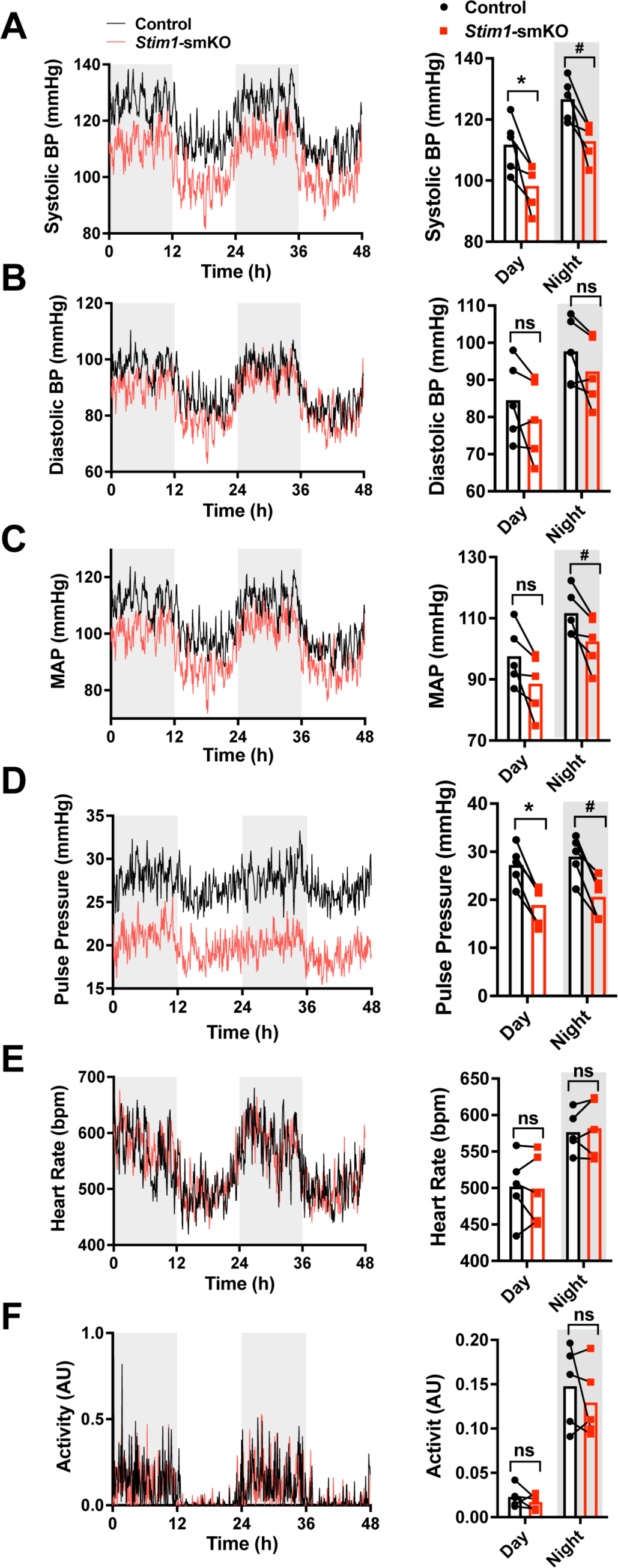
*Stim1*-smKO mice are hypotensive. **(A)** Systolic BP (mmHg) over 48 hours in conscious, radio telemeter-implanted *Stim1*-smKO mice before (control) and after (*Stim1*-smKO) tamoxifen injection. Shaded regions depict night cycles (n = 5 for both groups; *P < 0.05 vs. control day, ^#^P < 0.05 vs. control night, paired t-test). **(B)** Diastolic BP measurements for *Stim1*-smKO mice before and after tamoxifen injection (n = 5 for both groups, paired t-test). ns, not significant. **(C)** MAP for *Stim1*-smKO mice before and after tamoxifen injection (n = 5 for both groups, ^#^P < 0.05 vs. control night, paired t-test) ns, not significant. **(D)** Pulse pressure for *Stim1*-smKO mice before and after tamoxifen injection (n = 5 for both groups; *P < 0.05 vs. control day, ^#^P < 0.05 vs. control night, paired t-test). **(E)** HR for *Stim1*-smKO mice before and after tamoxifen injection (n = 5 for both groups, paired t-test) ns, not significant. **(F)** Locomotor activity (arbitrary units) for *Stim1*-smKO mice before and after tamoxifen injection (n = 5 for both groups, paired t-test) ns, not significant. Forty-eight-hour recordings are shown as means; bar graphs are shown as means ± SEM.

## Discussion

Junctional membrane complexes formed by close interactions of the ER/SR with the PM are critical signaling hubs that regulate homeostatic and adaptive processes in nearly every cell type. The canonical function of STIM1 is to enable SOCE via Orai channels, but mounting evidence suggests that the protein has additional, SOCE-independent functions. Here we show that STIM1 is crucial for fostering SR-PM junctions and functional coupling between SR and PM ion channels that control VSMC contractility. In support of this concept, we found that the number and sizes of SR/PM coupling sites were significantly reduced in VSMCs from *Stim1-*smKO mice. *Stim1* knockout also altered the nanoscale architecture of ion channels in Ca^2+^-signaling complexes, transformed the properties of Ca^2+^ sparks, and diminished BK and TRPM4 channel activity under physiological recording conditions. Resistance arteries isolated from *Stim1-*smKO mice exhibited blunted responses to vasoconstrictor stimuli, and animals became hypotensive following acute knockout of *Stim1* in smooth muscle. Collectively, these findings demonstrate that STIM1 maintains stable peripheral coupling between the SR and PM of contractile VSMCs in a manner that is independent of Ca^2+^ store depletion and SOCE. Loss of peripheral coupling in VSMCs following *Stim1* knockout has profound consequences, disrupting arterial function and BP regulation.

The SR-PM signaling domains of VSMCs are less orderly compared with those in cardiac and skeletal muscle cells and remain incompletely characterized. SR-PM junctions within the transverse (T) tubules of cardiomyocytes and skeletal muscle cells have regular, repeating structures that are formed, in part, by cytoskeletal elements and proteins of the junctophilin (64–66) and triadin (67, 68) families. In VSMCs, which lack T-tubules, SR-PM interactions occur at peripheral coupling sites that form throughout the periphery with no apparent pattern of distribution. Our research team has previously identified vital roles for microtubule networks (52) and junctophilin 2 (JPH2) (49) in the formation of peripheral coupling sites in VSMCs. Here, we found that knockout of *Stim1* in VSMCs reduced the number and sizes of SR-PM colocalization sites, demonstrating that STIM1 is necessary for the formation of stable SR-PM junctions in VSMCs with intact SR Ca^2+^ stores. Why is STIM1 active under these conditions? A simple explanation is that resting SR [Ca^2+^] in fully differentiated, contractile VSMCs is sufficiently low to trigger constitutive activation of STIM1. This concept is supported by a report by Luik et al. (69), who showed that the half-maximal concentration (K1/2) of ER Ca^2+^ for the activation of ICRAC (Ca^2+^release-activated Ca^2+^ current) in Jurkat T cells is 169 μM and the K1/2 for redistribution of STIM1 to the PM is 187 μM. These data are in close agreement with another study, which reported that the K1/2 of ER Ca^2+^ for redistribution of STIM1 in HeLa cells was 210 μM and that for maximum redistribution was 150 μM (70). Few studies have reported measurements of SR [Ca^2+^] in native, contractile VSMCs. Using the low-affinity ratiometric Ca^2+^ indicator, mag-fura-2, one well-controlled study estimated that resting SR [Ca^2+^] in contractile SMCs was ∼110 μm (4). Under these conditions, STIM1 is expected to be in a fully active configuration. It is also possible that regional SR [Ca^2+^] levels near active Ca^2+^-release sites (RyRs and IP3R) are lower than global SR [Ca^2+^], which could further stimulate STIM1 activity at these sites and reinforce junctional coupling. Thus, we put forward the concept that STIM1 is in an active state in quiescent contractile smooth muscle and is necessary for the formation of PM-SR junctional membrane contacts vital for contractile function. Our data further imply that, as VSMCs transition to a proliferative phenotype during the development of disease states associated with vascular remodeling, SR Ca^2+^ levels increase, leading to STIM1 inactivation, loss of peripheral coupling, and acquisition of SOCE activity (71).

Ion channel proteins in the membranes of excitable cells form discreet clusters whose sizes are exponentially distributed, a phenomenon that has been suggested to occur through stochastic self-assembly (72). Here, we found that acute knockout of STIM1 in VSMCs reduced the mean sizes of BK, TRPM4, and IP3R protein clusters and slightly increased the mean size of RyR2 protein clusters. According to the stochastic model proposed by Sato et al. (72), the steady-state size of membrane protein clusters is limited by the probability of removal from the PM through recycling or degradation processes, with larger clusters having a higher likelihood of removal. Thus, the smaller size of BK, TRPM4, and IP3R clusters following STIM1 knockout is likely a consequence of an increase in the rate of channel removal from the membrane. Accordingly, we propose that STIM1 increases the dwell time of BK, TRPM4, and IP3Rs proteins in the membrane, allowing larger clusters to form. This could occur through direct protein-protein interactions or via an indirect mechanism. Previous studies have provided evidence of direct interactions between STIM1 and IP3Rs (73, 74), but interactions between STIM1 and BK or TRPM4 have not been reported. It is also possible that intact SR-PM junctions partially protect membrane proteins from endocytic and/or recycling cascades, allowing larger clusters to form before they are removed. RyR2 cluster size was very slightly increased following STIM1 knockout, possibly reflecting the slow turnover rate of these massive proteins. Interestingly, we previously found that disruption of peripheral coupling in VSMCs by depolymerizing microtubules (52) or through morpholino-based knockdown of JPH2 (49) did not alter BK channel protein cluster sizes. This lack of an effect on cluster size could be attributable to the acute methods used to interrupt peripheral coupling in these previous studies or to specific properties of SR-PM junctional sites maintained by STIM1.

Knockout of *Stim1* in VSMCs significantly impacted Ca^2+^ signaling, ion channel activity, vascular contractility, and the regulation of BP. We purport that all of these outcomes result from nanoscale disruptions in cellular architecture. The compromised structural integrity of subcellular Ca^2+^ signaling microdomains formed by interactions of the PM and SR likely accounts for the more extensive spatial spread of Ca^2+^ sparks and prolonged clearance by SERCA pumps, the PMCA and Na^+^/Ca^2+^ exchangers, which extend the decay rate and duration of Ca^2+^ signals. Decreased nanoscale colocalization of BK with RyR2 and TRPM4 with IP3Rs manifested as diminished Ca^2+^-dependent activity of BK and TRPM4 channels (STOCs and TICCs), reflecting a loss in the functional coupling of Ca^2+^-release sites on the SR and ion channels on the PM. The smaller sizes of BK and TRPM4 protein clusters on the PM following *Stim1* knockdown may also reduce BK and TRPM4 channel currents. At the intact blood vessel level, the diminished TRPM4 and BK channel activity resulted in impaired contractility in response to physiological stimuli. This finding is interesting because our prior studies investigating the role of microtubular structures (52) and JPH2 (49) in maintaining peripheral coupling in VSMCs showed that disruption of PM-SR interactions caused cerebral arteries to become hypercontractile. In these studies, arterial hypercontractility resulted from interruption of the BK-RyR2 signaling pathway, which hyperpolarizes the VSMC membrane and balances the depolarizing and contractile influences of the TRPM4-IP3R cascade. *Stim1* knockout, in contrast, affected both pathways, indicating that STIM1 influences peripheral coupling in a manner that differs from that of the microtubule network and JPH2 and further suggesting heterogeneity in the formation of junctional membrane complexes in VSMCs. Diminished arterial contractility following *Stim1* knockout resulted in a drop in arterial BP, probably owing to a decrease in total peripheral resistance. This finding differs from previous reports by other groups showing that, although myogenic tone and phenylephrine-induced vasoconstriction was blunted in mesenteric arteries from a constitutive SMC-specific STIM1-knockout model, resting BP was not affected in this model (75–77). This difference is likely due to elevated levels of circulating catecholamines, which increase HR and cardiac output and thereby compensate for diminished vascular resistance (77).

In summary, our data demonstrate a vital role for STIM1 in the formation and maintenance of critical Ca^2+^-signaling microdomains in contractile VSMCs that is independent of SR Ca^2+^ store depletion. Disruptions in cellular architecture at the nanoscale level associated with the loss of STIM1 resulted in arterial dysfunction and impaired BP regulation, highlighting the essential nature of SR-PM junctions in cardiovascular control.

## Methods

### Animals

All animal studies were performed in accordance with guidelines of the Institutional Animal Care and Use Committee (IACUC) of the University of Nevada, Reno. Mice were housed in cages on a 12-hour/12-hour day-night cycle with ad libitum access to food (standard chow) and water. All transgenic mouse strains were obtained from The Jackson Laboratory (Bar Harbor, ME, USA). Mice with *loxP* sites flanking exon 2 of the *Stim1* gene (*Stim1^fl/f^*^l^ mice) were crossed with myosin heavy chain 11 (*Myh11*)-*cre*/ERT2 mice (47, 48), generating *Myh11-Cre-Stim1^fl/wt^* mice. Heterozygous *Myh11-Cre-Stim1^fl/wt^* mice were then intercrossed, yielding tamoxifen-inducible, SMC-specific *Stim1-*knockout mice (*Myh11*-*Cre*-*Stim1^fl/fl^*).

### Induction of STIM1 knockout

Male *Myh11*-*Cre*-positive *Stim1*-floxed mice (*Myh11*-*Cre*-*Stim1^fl/fl^*) were intraperitoneally injected at 4–6 weeks of age with 100 μL of a 10 mg/mL tamoxifen solution once daily for 5 days. Mice were used for experiments 1 week after the final injection. Littermate *Myh11*-*Cre*-*Stim1^fl/fl^* mice injected with the vehicle for tamoxifen (sunflower oil) were used as controls for all experiments.

### Wes capillary electrophoresis

Tissues isolated from mice were homogenized in ice-cold RIPA buffer (25 mM Tris pH 7.6, 150 mM NaCl, 1% Igepal CA-630, 1% sodium deoxycholate, 0.1% SDS) with protease inhibitor cocktail (Cell Biolabs, Inc., San Diego, CA) using a mechanical homogenizer followed by sonication. The resulting homogenate was centrifuged at 14,000 rpm for 20 minutes at 4°C, and the supernatant containing proteins was collected. Protein concentration was quantified with a BCA protein assay kit (Thermo Scientific, Waltham, MA) by absorbance spectroscopy using a 96-well plate reader. Proteins were then resolved by capillary electrophoresis using the Wes system (ProteinSimple, San Jose, CA, USA) and probed with an anti-STIM1 primary antibody (S6072; Sigma-Aldrich, St. Louis, MO, USA). Bands were analyzed using Compass for SW (ProteinSimple).

### SMC isolation

Mice were euthanized by decapitation and exsanguination under isoflurane anesthesia. Cerebral pial arteries were isolated carefully in ice-cold Mg^2+^-containing physiological salt solution (Mg^2+^-PSS; 5 mM KCl, 140 mM NaCl, 2 mM MgCl2, 10 mM HEPES, and 10 mM glucose; pH 7.4, adjusted with NaOH) and then incubated in an enzyme cocktail containing 1 mg/mL papain (Worthington Biochemical Corp., Lakewood, NJ, USA), 1 mg/mL dithiothreitol (DTT; Sigma-Aldrich), and 10 mg/mL bovine serum albumin (BSA; Sigma-Aldrich) for 12 minutes at 37°C. The arteries were then washed three times with Mg^2+^-PSS and incubated in 1 mg/mL collagenase type II (Worthington) in Mg^2+^-PSS for 14 minutes. The arteries were washed three times with Mg^2+^-PSS and then dissociated into single cells by triturating with a fire-polished glass Pasteur pipette.

### Visualization of PM-SR colocalization sites using SIM

Cerebral pial artery SMCs were allowed to adhere onto poly-L-lysine-coated round coverslips (5 mm diameter) during a 30-minute incubation at 37°C with the SR stain, ER-Tracker Green (Thermo Fisher Scientific; diluted 1:1000 in Mg^2+^-PSS). After incubation, ER-Tracker Green was removed, and the PM stain Cell-Mask Deep Red (Thermo Fisher Scientific; diluted 1:1000 in Mg^2+^-PSS) was added, and cells were incubated for 5 minutes at 37°C. Cell-Mask Deep Red was then removed and cells were washed with Mg^2+^-PSS and imaged using a lattice light-sheet microscope (LLSM; Intelligent Imaging Innovations, Inc., Denver, CO) (78). Coverslips with stained cells were mounted onto a sample holder and placed in the LLSM bath, immersed in Mg^2+^-PSS. Imaging was performed in SR-SIM mode, set to 100-ms exposures. For each cell, 200 Z-steps were collected at a step size of 0.25 µm. Imaging was limited to no more than 30 minutes for each coverslip to prevent artifacts caused by internalization of the plasma membrane dye. Surface reconstruction and colocalization analyses of PM and SR were performed using Imaris (Bitplane, Zurich, Switzerland) image analysis software. The Surface-Surface coloc plugin was used to visualize areas of the PM and SR that colocalized to form coupling sites.

### GSDIM superresolution microscopy

Ground state depletion microscopy followed by individual molecule return (GSDIM) was performed as described previously (49, 51, 52, 55, 56). Briefly, freshly isolated cerebral pial artery SMCs were allowed to adhere onto poly-L-lysine coated glass coverslips for 30 minutes. The cells were then fixed for 20 minutes with 2% paraformaldehyde, quenched with 0.4 mg/mL NaBH4, and permeabilized with 0.1% Triton X-100. Thereafter, cells were blocked with 50% SEABLOCK blocking buffer (Thermo Fisher Scientific, Waltham, MA) for 2 hours and incubated overnight at 4°C with primary antibody (Anti-STIM1 – (4916) Cell Signaling Technologies, Danvers, MA; Anti-BKα1 – (APC-021) Alomone Labs, Jerusalem, Israel; Anti-RyR2 – (MA3-916) Thermo Fisher Scientific, Waltham, MA; Anti-TRPM4 – (ABIN572220) antibodies-online.com, Limerick, PA; Anti-IP3R – (ab5804) Abcam, Cambridge, UK) diluted in PBS containing 20% SEABLOCK, 1% BSA, and 0.05% Triton X-100. Cells were washed three times with 1X PBS after each step. After overnight incubation, unbound primary antibody was removed by washing four times with 20% SEABLOCK, after which cells were incubated with secondary antibodies (Alexa-Fluor 647–or Alexa-Fluor 568–conjugated goat anti-rabbit, goat anti-mouse, donkey anti-goat or donkey anti-rabbit as appropriate) at room temperature for 2 hours in the dark. After washing with 1X PBS, coverslip-plated cells were mounted onto glass depression slides in a thiol-based photo-switching imaging buffer consisting of 50 mM Tris/10 mM NaCl (pH 8), 10% glucose, 10 mM mercaptoethylamine, 0.48 mg/mL glucose oxidase, and 58 μg/mL catalase. Coverslips were sealed to depression slides with Twinsil dental glue (Picodent, Wipperfurth, Germany) to exclude oxygen and prevent rapid oxidation of the imaging buffer. Superresolution images were acquired in epifluorescence mode using a GSDIM imaging system (Leica, Wetzlar, Germany) equipped with an oil-immersion 160× HCX Plan-Apochromat (NA 1.47) objective, an electron-multiplying charge-coupled device camera (EMCCD; iXon3 897; Andor Technology, Belfast, UK), and 500-mW, 532- and 642-nm laser lines. Localization maps were constructed from images acquired at 100 Hz for 25,000 frames using Leica LAX software. Post-acquisition image analyses of cluster size distribution were performed using binary masks of images in NIH ImageJ software. Object-based analysis was used to establish colocalization of proteins of interest in superresolution localization maps.

### Patch-clamp electrophysiology

Freshly isolated cerebral artery SMCs were transferred to the recording chamber and allowed to adhere to glass coverslips at room temperature for 20 minutes. Recording electrodes (3–4 MΩ) were pulled on a model P-87 micropipette puller (Sutter Instruments, Novado, CA, USA) and polished using a MF-830 MicroForge (Narishige Scientific Instruments Laboratories, Long Island, NY, USA). Spontaneous transient outward currents (STOCs) and transient inward cation currents (TICCs) were recorded in Ca^2+^-containing PSS (134 mM NaCl, 6 mM KCl, 1 mM MgCl2, 2 mM CaCl2, 10 mM HEPES, and 10 mM glucose; pH 7.4, adjusted with NaOH). The patch pipette solution contained 110 mM K-aspartate, 1 mM MgCl2, 30 mM KCl, 10 mM NaCl, 10 mM HEPES, and 5 μM EGTA (pH 7.2, adjusted with NaOH). Amphotericin B (200 µM), prepared on the day of the experiment, was included in the pipette solution to perforate the membrane. For all experiments, currents were recorded using an Axopatch 200B amplifier equipped with an Axon CV 203BU headstage (Molecular Devices). Currents were filtered at 1 kHz, digitized at 40 kHz, and stored for subsequent analysis. Clampex and Clampfit (version 10.2; Molecular Devices) were used for data acquisition and analysis, respectively. For STOCs, cells were clamped at a membrane potential manually spanning a range from −60 mV to 0 mV. STOCs were defined as events > 10 pA, and their frequency was calculated by dividing the number of events by the time between the first and last event. Whole-cell K^+^ currents were recorded using a step protocol (−100 to +100 mV in 20 mV steps for 500 ms) from a holding potential of −80 mV. Whole-cell BK currents were calculated by current subtraction following administration of the selective BK channel blocker paxilline (1 μM). Current–voltage (I–V) plots were generated using currents averaged over the last 50 ms of each voltage step. The bathing solution contained 134 mM NaCl, 6 mM KCl, 10 mM HEPES, 10 mM glucose, 2 mM CaCl2, and 1 mM MgCl2; pH 7.4 (NaOH). The pipette solution contained 140 mM KCl, 1.9 mM MgCl2, 75 μM Ca^2+^, 10 mM HEPES, 0.1 mM EGTA, and 2 mM Na2ATP; pH 7.2 (KOH).

TICCs, induced by membrane stretch delivered by applying negative pressure (20 mmHg) through the recording electrode using a Clampex controlled pressure clamp HSPC-1 device (ALA Scientific Instruments Inc., Farmingdale, NY, USA), were recorded from cells clamped at a membrane potential of −70 mV. TICC activity was calculated as the sum of the open channel probability (*NP*o) of multiple 1.75-pA open states (5). Conventional whole-cell TRPM4 currents were recorded using ramp protocol consisting of a 400 ms increasing ramp from −100 to +100 mV ending with 300 ms step at +100 mV from a holding potential of −60 mV. A new ramp was applied every 2 s. TRPM4 whole-cell currents were recorded in a bath solution consisting of (in mM): 156 NaCl, 1.5 CaCl2, 10 glucose, 10 HEPES and 10 TEA-Cl; pH 7.4 (NaOH). The patch pipette solution contained (in mM): 156 CsCl, 8 NaCl, 1 MgCl2 10 mM HEPES; pH 7.4 (NaOH) and 200 µM free [Ca^2+^], adjusted with appropriate amount of CaCl2 and EGTA as calculated using Max-Chelator software.

### Quantitative droplet digital PCR

Total RNA was extracted from arteries by homogenization in TRIzol reagent (Invitrogen, Carlsbad, CA), followed by purification using a Direct-zol RNA microprep kit (Zymo Research, Irvine, CA), DNase I treatment (Thermo Fisher Scientific), and reverse transcription into cDNA using qScript cDNA Supermix (Quanta Biosciences, Gaithersburg, MD). Quantitative droplet digital PCR (ddPCR) was performed using QX200 ddPCR EvaGreen Supermix (Bio-Rad, Hercules, CA), custom-designed primers (Supplementary table S1), and cDNA templates. Generated droplet emulsions were amplified using a C1000 Touch Thermal Cycler (Bio-Rad), and the fluorescence intensity of individual droplets was measured using a QX200 Droplet Reader (Bio-Rad) running QuantaSoft (version 1.7.4; Bio-Rad). Analysis was performed using QuantaSoft Analysis Pro (version 1.0.596; Bio-Rad).

### Ca^2+^ imaging

A liquid suspension (∼0.2 mL) of freshly isolated VSMCs was placed in a recording chamber (RC-26GLP, Warner Instruments, Hamden, CT, USA) and allowed to adhere to glass coverslips for 20 minutes at room temperature. VSMCs were then loaded with the Ca^2+^-sensitive fluorophore, Fluo-4 AM (1 µM; Molecular Probes), in the dark for 20 minutes at room temperature in Mg^2+^-PSS. Cells were subsequently washed three times with Ca^2+^-containing PSS and incubated at room temperature for 20 minutes in the dark to allow sufficient time for Fluo-4 de-esterification. Images were acquired using an iXon 897 EMCCD camera (Andor; 16 x 16 µm pixel size) coupled to a spinning-disk confocal head (CSU-X1; Yokogawa), with a 100x oil-immersion objective (Olympus; NA 1.45) at an acquisition rate of 33 frames per second (fps). Custom software provided by Dr. Adrian D. Bonev (University of Vermont) was used to analyze the properties of Ca^2+^ sparks. The threshold for Ca^2+^ spark detection was defined as local increases in fluorescence *≥* 0.2 ΔF/F0.

### Pressure myography

Cerebral pial and 3^rd^ order mesenteric arteries were carefully isolated in ice-cold Mg^2+^-PSS. Each artery was then cannulated and mounted in an arteriography chamber and superfused with oxygenated (21% O2/6% CO2/73% N2) Ca^2+^-PSS (119 mM NaCl2, 4.7 mM KCl, 21 mM NaCO3, 1.18 mM KH2PO4, 1.17 mM MgSO4, 0.026 mM EDTA, 1.8 mM CaCl2, and 4 mM glucose) at 37°C and allowed to stabilize for 15 minutes. Each artery was then pressurized to 110 mmHg using a pressure servo controller (Living Systems instruments, St. Albans City, VT, USA). Any kinks or bends were gently straightened out, the pressure was reduced to 5 mmHg, and the artery was allowed to stabilize for 15 minutes. The viability of each artery was assessed by measuring the response to high extracellular [K^+^] PSS (made isotonic by adjusting the [NaCl], 60 mM KCl, 63.7 mM NaCl). Arteries that contracted less than 10% were excluded from further investigation.

Myogenic tone was assessed by raising the intraluminal pressure from 5 mmHg to 140 mmHg in 20-mmHg increments, with the artery maintained at each pressure increment for 5 minutes (active response). The artery was then superfused for 15 minutes at 5 mmHg intraluminal pressure with Ca^2+^-free PSS supplemented with EGTA (2 mM) and the voltage-dependent Ca^2+^ channel blocker diltiazem (10 μM), followed by application of pressure increments from 5 mmHg to 140 mmHg (passive response). The artery lumen diameter was recorded using edge-detection software (IonOptix, Westwood, MA, USA). Myogenic reactivity at each intraluminal pressure was calculated as [1 – (Active diameter/Passive Diameter)] *×* 100.

In separate arteries, the contractile response to the thromboxane A2 receptor agonist U46619 and α1-adrenergic receptor agonist phenylephrine was assessed in cerebral and mesenteric arteries, respectively. Arteries were pressurized to 20 mmHg to prevent the development of myogenic tone. Cumulative concentration response curves were produced through the addition of U46619 (0.01–1000 nM) or phenylephrine (0.001–100 µM) to the superfusing bath solution. Arteries were mantained at each concentration for 5 minutes or until a steady-state diameter was reached, before addition of the next concentration. Following the addition of the final concentration, arteries were bathed in Ca^2+^-free PSS to obtain the passive diameter. Contraction was calculated at each concentration as vasoconstriction (%) = [(lumen diameter at constriction − lumen diameter at baseline)/passive lumen diameter] × 100.

### In vivo radiotelemetry

*Stim1*-smKO mice were initially anesthetized using 4–5% isoflurane carried in 100% O2 (flushed at 1 L/min), after which anesthesia was maintained by adjusting isoflurane to 1.5–2%; preoperative analgesia was provided by subcutaneous injection of 50 µg/kg buprenorphine (ZooPharm, Windsor, CO, USA). The neck was shaved and then sterilized with iodine. Under aseptic conditions, an incision (∼1 cm) was made to separate the oblique and tracheal muscles and expose the left common carotid artery. The catheter of a radio telemetry transmitter (PA-C10; Data Science International, Harvard Bioscience, Inc., Minneapolis, MN, USA) was surgically implanted in the left common carotid artery and secured using non-absorbable silk suture threads. The body of the transmitter was embedded in a subcutaneous skin pocket under the right arm. After a 14-day recovery period, baseline BP, HR, and locomotor activity were recorded in conscious mice for 48 hours using Ponemah 6.4 software (Data Science International). Parameters were measured for 20 seconds every 5 minutes. Mice were then injected with either vehicle or tamoxifen using the protocol described above; after 7 days following the final injection, baseline BP readings, HR, and locomotor activity were re-recorded in conscious mice for 48 hours

### Chemicals

All chemicals used were obtained from Sigma-Aldrich (St. Louis, MO, USA) unless specified otherwise.

### Statistical analysis

All data are expressed as means ± standard error of the mean (SEM) unless specified otherwise. Statistical analyses were performed using either unpaired Student’s t-test or analysis of variance (ANOVA) as appropriate, and a P-value < 0.05 was considered statistically significant. GraphPad Prism v8.2 (GraphPad Software, Inc., USA) was used for statistical analyses and graphical presentations.

## Author Contributions

S.E., A.L.G., and M.T. conceived and initiated the project. S.E. supervised the project and designed experiments. V.K. performed GSDIM and SIM superresolution imaging and Wes protein-detection experiments. S.A. performed patch-clamp electrophysiology experiments. C.S.G. conducted and analyzed Ca^2+^ spark recordings. P.T. performed pressure myography and *in vivo* BP recording studies. E.Y. performed RT-ddPCR experiments. M.J. helped with the development of transgenic animal models and immunolabeling protocols. A.L.G. performed experiments demonstrating the feasibility of the project. V.K., S.A., P.T., E.Y., M.G.A., C.S.G., and S.E. analyzed the data. V.K. and S.E. wrote the manuscript and prepared the figures. V.K., S.A., A.L.G., M.T., and S.E. revised the manuscript.

## Supplementary Figures

**Supplementary figure 1:**
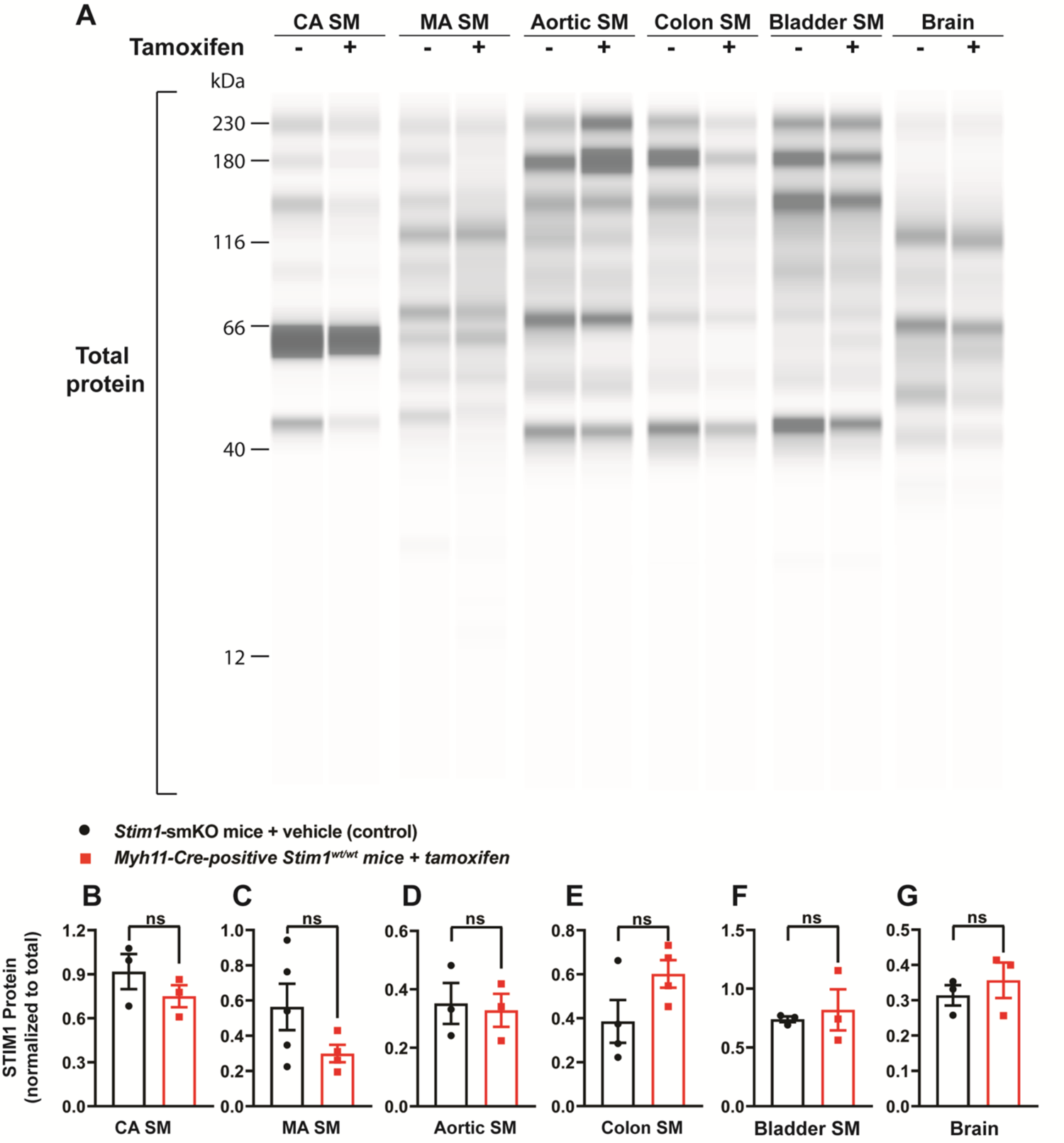
**(A)** Representative Wes blot showing total protein levels in cerebral artery smooth muscle (CA SM), mesenteric artery smooth muscle (MA SM), aortic smooth muscle, colonic smooth muscle, bladder smooth muscle, and whole-brain tissues isolated from vehicle- and tamoxifen-injected *Stim1-*smKO mice**. (B–G)** Summary data showing STIM1 protein expression normalized to total protein levels in cerebral artery smooth muscle **(B)**, mesenteric artery smooth muscle **(C)**, aortic smooth muscle **(D)**, colonic smooth muscle **(E)**, bladder smooth muscle **(F)**, and brain **(G)** tissues isolated from vehicle-injected *Stim1-*smKO mice (control) and tamoxifen-injected *Stim1-*smKO mice (n = 3–5 mice/group, unpaired t-test) ns, not significant.

**Supplementary figure 2:**
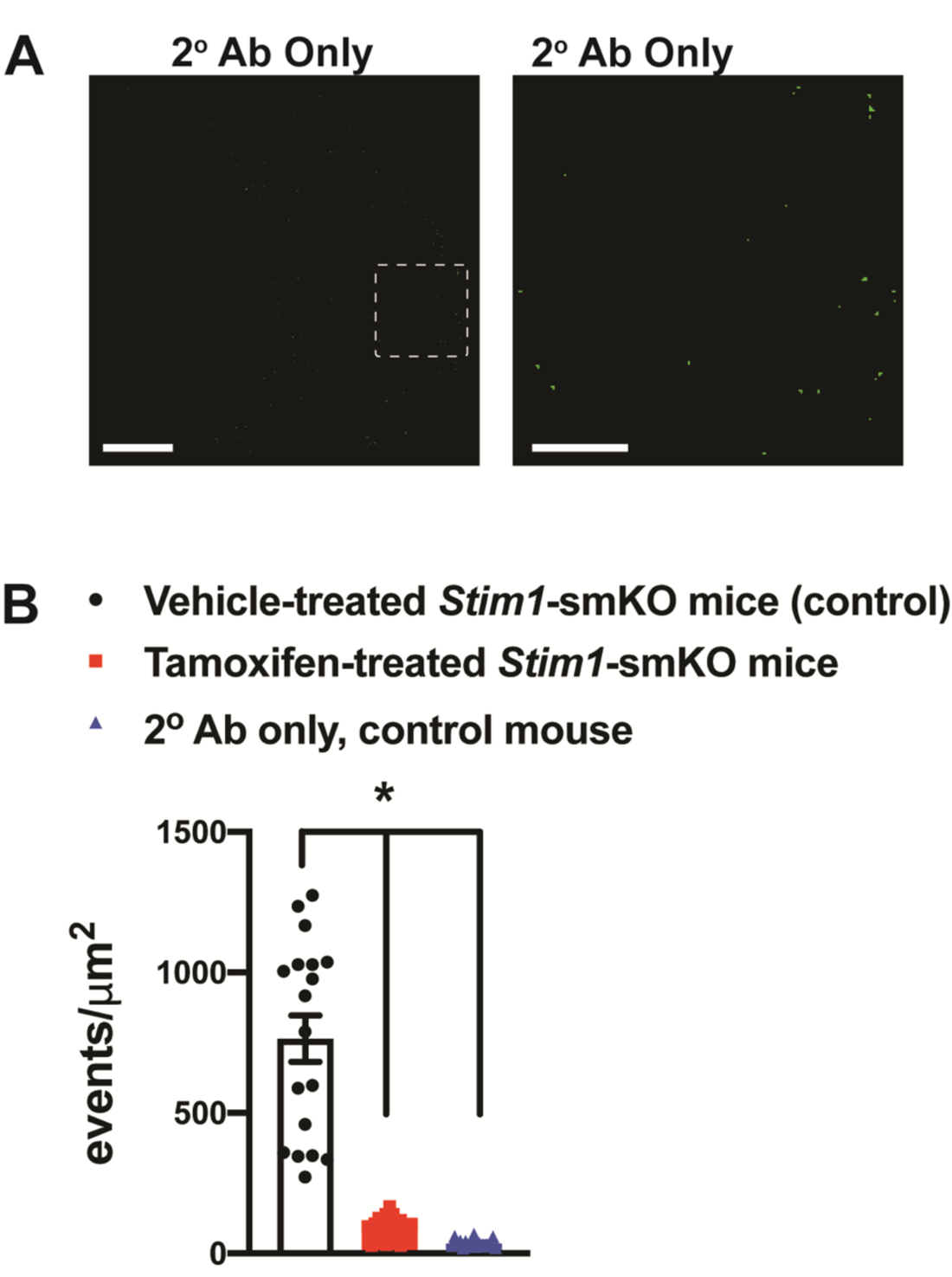
**(A)** Representative superresolution localization maps of an isolated cerebral artery SMC from a vehicle-injected (control) *Stim1-*smKO mouse immunolabeled with only the secondary antibody (2° Ab) used to detect STIM1 (goat anti-rabbit Alexa Fluor 647) and imaged with GSDIM. Right panel: enlarged image of the area highlighted by the white square in the left panel. Scale bars: 3 μm (left panel) and 1 μm (right panel). **(B)** GSDIM events detected per unit area in cerebral artery SMCs isolated from vehicle- and tamoxifen-injected *Stim1-*smKO mice immunolabeled with anti-STIM1 antibody and cells isolated from vehicle-injected *Stim1-*smKO mice immunolabeled only with the secondary antibody (negative control). Data are presented as means ± SEM (n = 18 cells from 3 mice in each group; *P < 0.05 vs. control, one-way ANOVA).

**Supplementary figure 3:**
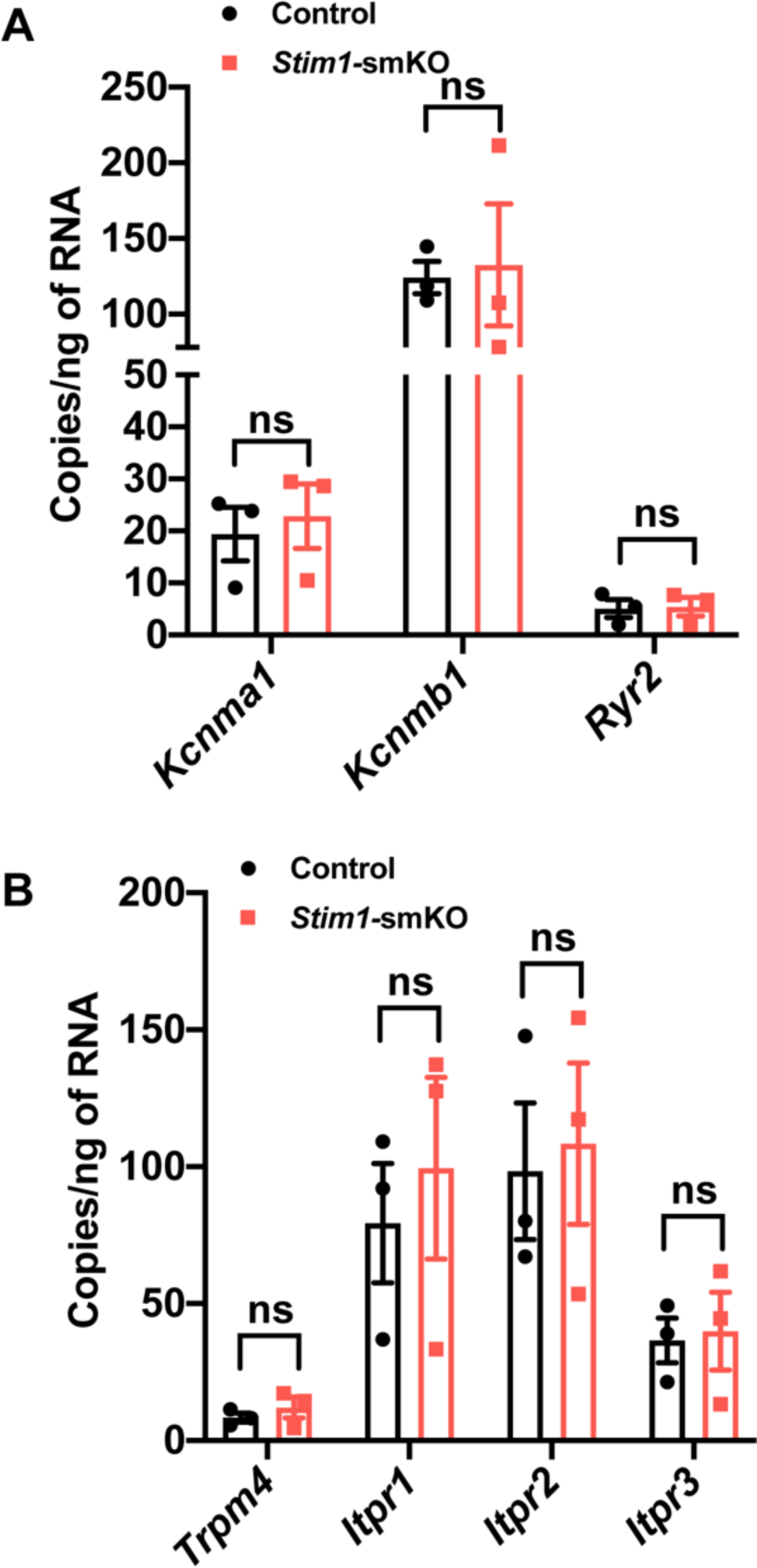
**(A)** mRNA expression levels (transcript copies/ng of RNA) of *Kcnma1* (BK*α*1), *Kcnmb1* (BK*β*1), and *Ryr2* (RyR2) in cerebral arteries from control and *Stim1*-smKO mice, determined by quantitative droplet digital PCR (ddPCR). **(B)** mRNA expression levels of *Trpm4* (TRPM4), *Itpr1* (IP3R1), *Itpr2* (IP3R2), and *Itpr3* (IP3R3) in cerebral arteries from control and *Stim1*-smKO mice. Data are presented as means ± SEM (n = 3 mice in each group, unpaired t-test). ns, not significant.

**Supplementary figure 4:**
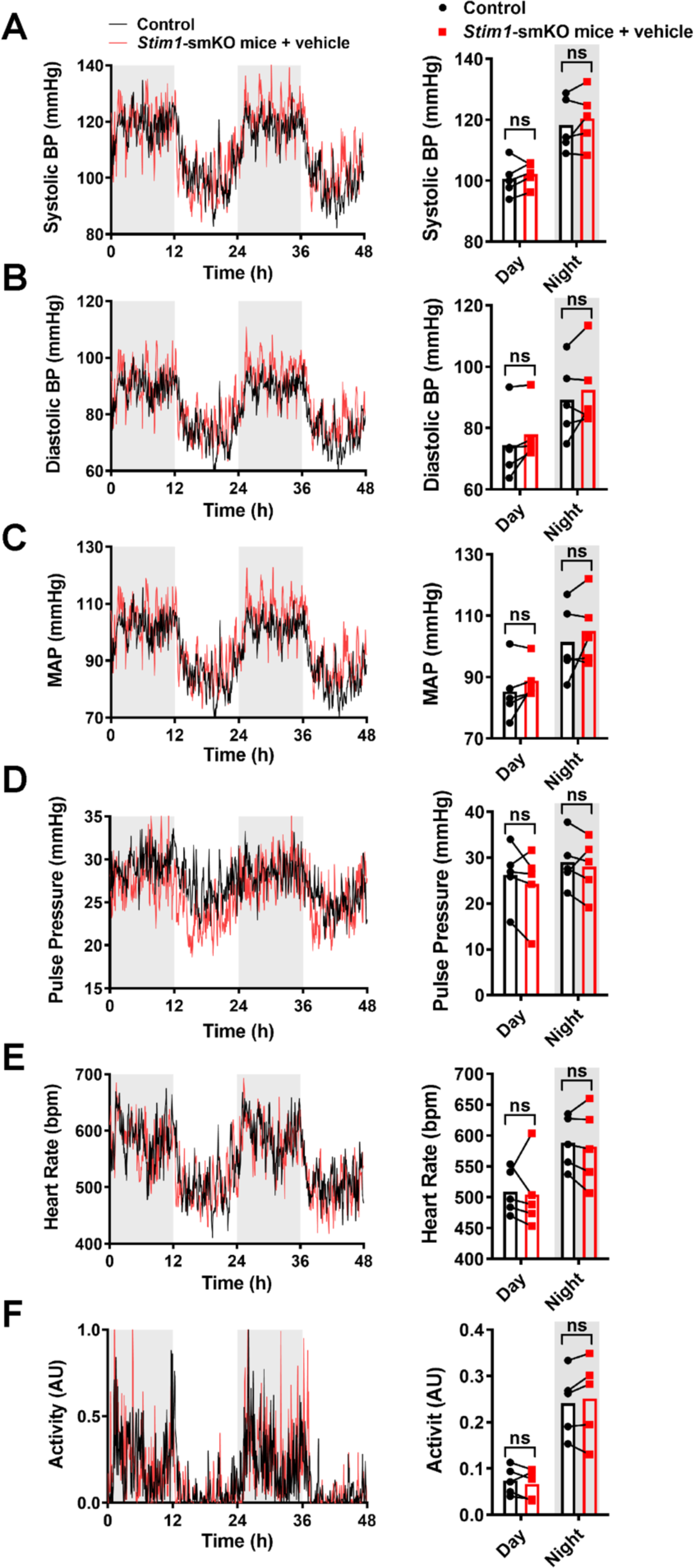
**(A)** Systolic BP (mmHg) over 48 hours in conscious, radio telemeter-implanted *Stim1*-smKO mice before (control) and after vehicle injection. Shaded regions depict night cycles (n = 5 for both groups). **(B)** Diastolic BP measurements for *Stim1*-smKO mice before and after vehicle injection (n = 5 for both groups). **(C)** MAP for *Stim1*-smKO mice before and after vehicle injection (n = 5 for both groups). **(D)** Pulse pressure for *Stim1*-smKO mice before and after vehicle injection (n = 5 for both groups). **(E)** HR for *Stim1*-smKO mice before and after vehicle injection (n = 5 for both groups). **(F)** Locomotor activity (arbitrary units [AU]) for *Stim1*-smKO mice before and after vehicle injection (n = 5 for both groups). Forty-eight-hour recordings are shown as means; bar graphs are shown as means ± SEM. There were no significant differences.

## Supplementary Table

**Table S1.**
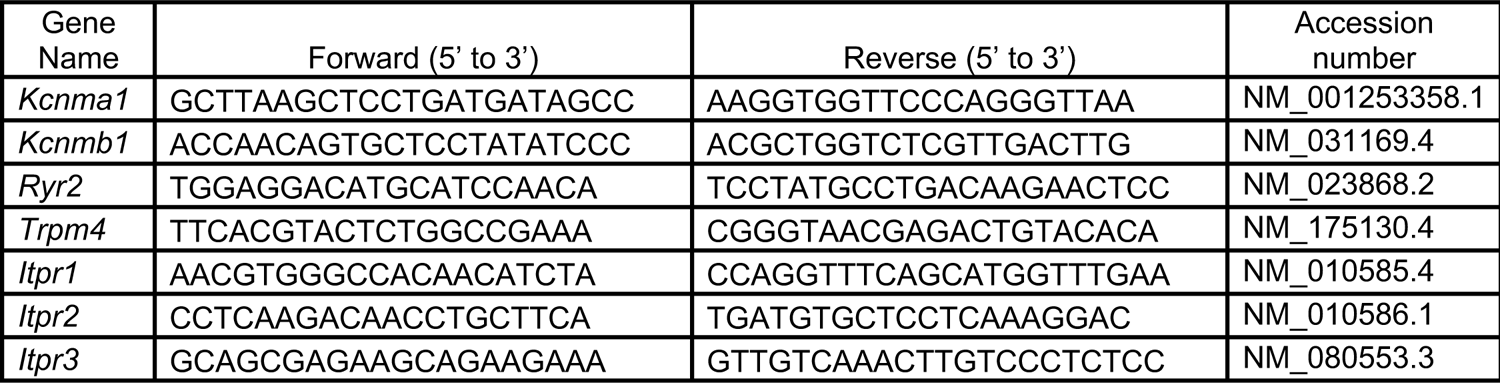
Forward and reverse primer sequences used for ddPCR experiments.

## Supplementary Movie Legends

**Supplementary movie 1:**

PM-SR interactions in a cerebral artery SMC isolated from a control mouse. Animated representation of a SIM image series that was reconstructed and rendered in 3D. The PM is shown in red and made transparent for better visualization; the SR is shown in green; and colocalized areas are shown in yellow.

**Supplementary movie 2:**

PM-SR interactions in a cerebral artery SMC isolated from a tamoxifen-injected *Stim1-*smKO mouse. Animated representation of a SIM image series that was reconstructed and rendered in 3D. The PM is shown in red and made transparent for better visualization; the SR is shown in green; and areas of colocalization are shown in yellow.

**Supplementary movie 3:**

Representative movie showing spontaneous Ca^2+^ sparks in a cerebral artery SMC isolated from a control mouse.

**Supplementary movie 4:**

Representative movie showing spontaneous Ca^2+^ sparks in a cerebral artery SMC isolated from a *Stim1-*smKO mouse.

